# Hbs and Rst adhesion molecules provide a regional code that regulates cell elimination during epithelial remodelling

**DOI:** 10.1101/2025.07.25.666765

**Authors:** Miguel Ferreira-Pinto, Mario Aguilar-Aragon, Christa Rhiner, Eduardo Moreno

## Abstract

Cell–cell interactions and mechanical forces are fundamental in shaping epithelial tissue architecture. For example, tissue compression promotes cell elimination by downregulating compaction-sensitive EGFR/ERK signalling. However, a subgroup of cells that are eliminated do not show EGFR/ERK downregulation, suggesting the involvement of additional mechanisms. Here, we conducted a multi-step RNAi screen in the *Drosophila* notum and identified 47 diverse regulators of epithelial remodelling. Among them, we focused on two membrane proteins, Hibris (Hbs) and Roughest (Rst), which mediate heterophilic cell adhesion. We show that Hbs and Rst exhibit region-specific expression patterns in the developing notum, showing elevated protein levels in zones of cell survival and reduced expression in regions undergoing cell pruning. Uniform knock-down of *hbs* and *rst* or homogenous overexpression of *hbs*, disrupts normal cell elimination during morphogenesis and results in adult tissue malformations. Remarkably, local suppression of Hbs and Rst in Hbs^high^ /Rst^high^ territories triggers ectopic cell elimination, indicating that a surface code defined by Hbs/Rst can instruct cell removal. We further demonstrate that Hbs, but not Rst, is regulated by compaction-sensitive EGFR signalling, positioning Hbs as a potential integrator of mechanical cues and cell property signals, possibly via its interaction with Rst. These findings uncover a novel adhesive code that shapes the thorax midline and potentially other organs.

## Introduction

All animals, regardless of their different sizes and shapes, develop based on fundamental rules that guide how tissues grow and organize. Epithelial tissues establish the body’s fundamental architecture, forming sheets and tubes that give rise to organs such as the gut, lungs, and skin. A finely tuned balance between cell division, cell death, cell migration and cell rearrangement ensures the precise tissue shape (Lecuit and Lenne, 2007). Mechanical forces drive crucial aspects of morphogenesis such as elongation or folding (Paci and Mao, 2021), cell-cell interactions (Eder, Aegerter and Basler, 2017; Brás-Pereira and Moreno, 2018; Matamoro-Vidal and Levayer, 2019) and adjusting the rate of cell death (Shraiman, 2005; Aegerter-Wilmsen *et al*., 2012; Marinari *et al*., 2012). Compression forces, in particular, have been shown to act as an integral-feedback mechanism, regulating the elimination of compacted suboptimal cells in epithelial co-cultures of Madin–Darby canine kidney (MDCK) cells, zebrafish epidermal fin edges and the *Drosophila* notum, an epithelium giving rise to adult dorsal thorax structure (Eisenhoffer *et al*., 2012; Marinari *et al*., 2012; Levayer, Dupont and Moreno, 2016; Wagstaff *et al*., 2016; Moreno *et al*., 2019).

In zebrafish, mechanical crowding engages the stretch-activating channel Piezo1 and sphingosine 1-phosphate signalling, which converges on Rho-kinase-dependent myosin contraction leading to cell extrusion (Eisenhoffer *et al*., 2012; Wagstaff *et al*., 2016). As a cell intrinsic factor, high levels of P53 have been shown to be sufficient to induce crowding hypersensitivity in MDCK co-cultures (Wagstaff *et al*., 2016; Bove *et al*., 2017). In *Drosophila*, cell elimination in the notum is also regulated by apoptotic components, requiring the induction of the pro-apoptotic factor *head involution defective* (*hid*) followed by caspase activation and microtubule disassembly, which precede cell extrusion (Levayer, Dupont and Moreno, 2016; Moreno *et al*., 2019; Villars *et al*., 2022).

Upstream of *hid* induction, the downregulation of compaction-induced EGFR/ERK survival signalling has been shown to be important for cell death at the notum midline (Moreno *et al*., 2019), however the nature of the mechano-sensing properties of EGFR during tissue compaction have remained unclear (Moreno *et al*., 2019; Crozet and Levayer, 2023). Interestingly, 25% of epithelial cells in the notum are being eliminated without signs of EGRF/ERK downregulation (Moreno *et al*., 2019). Moreover, Genes mediating fitness-dependent cell death such as *flower (fwe*) (Rhiner *et al*., 2010), *azot* (Merino *et al*., 2015) and JNK (Pereira *et al*., 2011; Yamamoto *et al*., 2017) did not show a role in controlling cell death in the notum (Levayer, Dupont and Moreno, 2016; Moreno *et al*., 2019). Together, these findings indicate that additional, yet unknown players must guide cell pruning in the notum, thereby ensuring correct epithelium remodelling.

Here, we used the well-established *Drosophila* pupal notum model to discover novel regulators of cell elimination. To this end, we conducted an unbiased “*in-silico*” genetic screen and identified 47 new modifiers with diverse molecular characteristics and subcellular localizations. Among them, the cell adhesion molecules Hibris (Hbs) and Roughest (Rst) were identified as top regulators of cell elimination. Interestingly, the two Immunoglobulin-like cell adhesion molecules (Ig-CAM) Hbs and Rst have been previously found to act as recognition module-proteins during the formation of the highly structured fly retina (Cagan and Ready, 1989; Bao and Cagan, 2005; Bao, 2014) and the patterning of sensory organs in the wing (Dworak *et al*., 2001; Takemura and Adachi-Yamada, 2011; Linneweber, Winking and Fischbach, 2015). In the retina, the heterophilic binding of Hbs and Rst maximizes the cell-cell contacts of a subset of interommatidial progenitor cells (IPS), while excess IPS with insufficient adhesion are eliminated (Bao and Cagan, 2005). Similarly, selective adhesion conferred by Hbs and Rst on different cell types are thought to mediate cell sorting into highly structured zigzag pattern of *Drosophila* wing margin hairs (Takemura and Adachi-Yamada, 2011). How these two cell adhesion molecules could regulate highly dynamic tissue movements in a developing epithelial sheet is not known and their link to mechanical forces remains unexplored.

In this study, we identify a new comprehensive set of candidate modulators of cell elimination in the notum. In particular, we uncover a novel role of Hbs and Rst in specifying an adhesive network during epithelial remodelling in the fly notum, which, when perturbed, leads to malformation in adult tissues. Our results show that Hbs and Rst relative levels are sufficient to instruct cell elimination or cell survival in the developing notum, functioning as a cell selection code that guides proper organization of epithelial cells. We find that the pattern of the Hbs/Rst adhesive landscape correlates with crowding and compaction areas and provide evidence that one component of Hbs/Rst surface code is modulated by compaction-sensitive EGFR/ERK signalling, previously shown to play a crucial role in cell selection at the notum midline. Our findings put forward a new framework, in which Hbs and Rst constitute new players of the regulatory network, potentially integrating both EGFR/ERK-dependent and independent signals to control epithelium architecture.

## Results and Discussion

### A multi-step screen to identify novel epithelial remodelling regulators

During pupal notum development, cells are eliminated via mechanical and non-mechanical inputs, ensuring the correct notum size and shape in adult flies (Figure 1A). As previously reported, cell elimination events are more frequent inside the midline region of the notum than outside, due to increased cell crowding at the midline (Figure 1A) (Levayer, Dupont and Moreno, 2016; Moreno *et al*., 2019). The notum midline cells give rise to the central section of the adult thorax (Figure 1B-1C). Therefore, the silencing of key cell death regulators in the developing notum, will directly affect the width of the adult midline region in the thorax. We tested this by silencing the pro-apoptotic gene *hid*, which resulted in an increased midline, whereas silencing of negative regulator EGFR led to a reduced midline (Figure 1D-1E). Based on these results, we concluded that the adult midline phenotype can serve as a reliable readout for identifying novel regulators of cell death.

**Figure 1.**
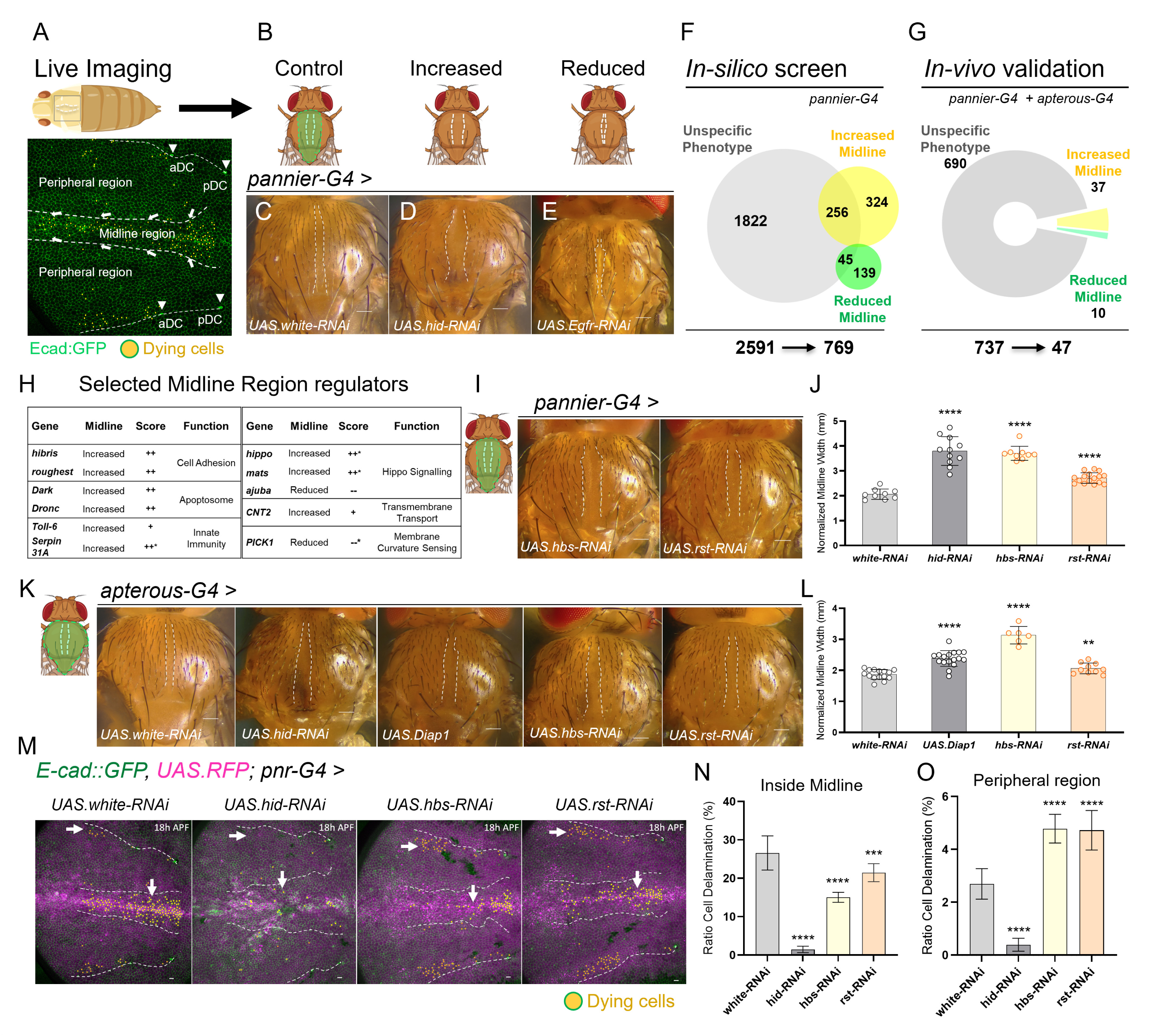
A multi-step screen identified Hbs and Rst as novel epithelial remodelling regulators. A) Schematic of the notum epithelium during pupal development. The central bristle row marks the midline (white dashed lines). Arrows indicate convergence cell flow towards the midline. Dying cells are shown in yellow. E-cadherin (green) labels cell membranes for tracking cell fate over time. The aDC/pDC macrochaetae delimit the peripheral region of the notum. B) Schematic illustrating different midline phenotypes (white dashed lines). The *pannier* domain is marked (green). (C–E) Adult notum images showing midlines of *pannier-Gal4* driven *white* RNAi (control), *hid* RNAi and *Egfr* RNAi. White dashed lines delineate the midline. Scale bars: 10 µm. F) Recovered midline phenotypes in the “*in-silico* screen” based on RNAi activation with *pannier-Gal4*. Overlapping areas indicate genes with both midline-specific and nonspecific phenotypes. G) Midline phenotypes assessed by *in-vivo* gene knock-down driven by *pannier* and *apterous-Gal4* notum drivers. H) Table highlighting selected midline regulators, validated *in-vivo*. Scores denote ++ strong midline increase; + intermediate midline increase; -- strong midline decrease; - intermediate midline decrease. Asterisks mark genes that caused mild notum defects outside the midline. I) Adult notum images showing increased midline width upon *hbs* and *rst* RNAi in the pannier domain. Scale bars: 10 µm. J) Quantification of normalised midline width (mm) upon gene downregulation using *pannier-Gal4*. Each dot represents one adult fly thorax. Statistical significance was assessed by unpaired t-test or Mann-Whitney against the control; ****p<0.0001. Bars represent standard deviation (SD). K) Adult notum images displaying midline phenotypes of control (*UAS.white* RNAi), *UAS.hid* RNAi, *UAS.Diap1*, *UAS.hbs* RNAi and *UAS.rst* RNAi driven by *apterous Gal-4*. L) Quantification of normalised midline width (mm) based on RNAi driven by *apterous-Gal4*. Each dot represents one adult fly thorax. Scale bars: 10 µm. Statistical significance was assessed by unpaired t-test or Mann-Whitney against the control; ****p<0.0001. M) Z-projections of live pupal nota at 18h APF (700min movies) with quantified cell death in control flies (*white* RNAi) and upon *hid* RNAi, *hbs* RNAi and *rst* RNAi. Yellow cells indicate future dying cells within the *pannier* domain (RFP, magenta). White arrows point to cell delamination events. Cell membrane is marked with *ubi-Ecad::GFP* (green). White dashed lines mark the midline. Peripheral regions analysed are delimited by the aDC and pDC bristles (white dashed lines). Scale bars: 10 µm. N-O) Quantification of cell delamination events (ratio) inside the midline region and in the periphery over 12 hours (700 min, from 18h to 30h APF). Bars represent the standard error of the mean (SEM) for each condition. Sample sizes: *white* RNAi (control): n = 4 nota, 1,531 cells (midline), 4,816 cells (outside); *hid* RNAi: n = 2 nota, 1,618 cells (midline), 3,148 cells (outside); *hbs* RNAi: n = 4 nota, 1,848 cells (midline), 5,118 cells (outside); *rst* RNAi: n = 4 nota, 2,071 cells (midline), 5,066 cells (outside). Statistical analysis was performed using Fisher’s exact test against the control; ****p < 0.0001).

To uncover new components mediating epithelial remodelling, we next devised a two-step genetic screen taking advantage of the midline phenotype. First, we conducted an *in-silico* screen, in which we analysed the width of the midline in a vast collection of adult fly notum images from the public Bristle Screen Database (BSD). The database provides notum images based on a genome-wide RNAi screen performed by the Knoblich Lab (Mummery-Widmer *et al*., 2009) (Supplementary Figure 1A). We quantified midline phenotypes in all available BSD pictures, corresponding to 4024 RNAi lines covering 2591 genes and categorized them into three different classes. Genes, for which the knock-down (RNAi) resulted in an altered midline - without causing major notum defects - were classified as either “Increased Midline” or “Reduced Midline” (Figure 1F and Supplementary Figure 1B). Genes, for which silencing affected the overall notum size or morphology, were excluded and grouped into the category “Unspecific” alongside genes that did not cause any significant midline change (Figure 1F and Supplementary Figure 1B). Although this approach may have led to the exclusion of cell death and remodelling regulators that, in addition, also cause disruption of other developmental processes, we obtained a total of 769 genes that caused a midline phenotype, which were retained for further analyses. In addition, we classified the strength of the observed midline phenotype, ranging from strong (++) to intermediate (+) while having, or not, minor notum defects (*) (Supplementary Figure 1B).

As different RNAi lines for the same gene occasionally produced dissimilar results, we included in the *in-silico* candidates all genes for which at least one RNAi line gave rise to a significant midline phenotype (769 genes) (Figure 1F). Precisely 95.8% of these genes, based on availability of RNAi lines, were next assessed in a robust “*in-vivo*” validation screen (Figure 1G), in which two independent RNAi lines were analysed per gene, in combination with two different notum drivers, *pannier-Gal4* and *apterous-Gal4* that drive gene expression in slightly different patterns in the pupal notum (Supplementary Figure 1C-1H). The strategy efficiently eliminated candidates that had been retained due to variable effects of the RNAi lines or inaccurate assessment of the midline phenotype based on BSD images. As a result, this composite large-scale screen with dual validation of candidates *in-vivo,* led to the identification of 47 high confidence candidates for epithelial remodelling at the pupal midline (Figure 1G and Supplementary Figure 1I). The midline phenotypes for all 47 candidates are shown in Supplementary Table 1 (scores) and Supplementary Figure 2 (images).

Among the 47 identified regulators of cell remodelling at the midline with 37 showing an increased midline and 10 a reduced midline, we found the apoptosome proteins Dark and Dronc, innate immunity components *toll-6* and *serpin31A*, Hippo signalling factors (*hippo*, *mats* and *ajuba*), the transmembrane transporter CNT2 and the cell adhesion molecules Hibris and Roughest (Figure 1H). The candidates span diverse functional groups, from transcription factors to cell tension regulators (Supplementary Figure 1J-1K). The pro-apoptotic genes were expected to be recovered in the approach as caspase activation were previously shown to control cell extrusion in the pupal notum (Levayer, Dupont and Moreno, 2016; Moreno *et al*., 2019). Beyond that, the screen uncovered many novel regulators with diverse functions whose role in epithelial cell selection and remodelling can be addressed in the future.

### Hibris and Roughest regulate cell pruning in the notum

Cell adhesion is an important quality to transmit mechanical forces across a tissue or mediate specific cell-cell recognition. We therefore focused on the two transmembrane proteins Hibris (Hbs) and Roughest (Rst) that localize to adherens junction. Hbs and Rst have been previously found to regulate cell rearrangement during *Drosophila* eye development via heterophilic binding (Bao and Cagan, 2005) and control sensory organs patterning in the wing (Dworak *et al*., 2001; Takemura and Adachi-Yamada, 2011; Linneweber, Winking and Fischbach, 2015). We found that RNAi of *hbs* and *rst* consistently resulted in an increased midline width for both tested RNAi lines and in conjunction with both *pannier* and *apterous* notum drivers (Figure 1I-1L). Suppression of Hibris caused a similarly increased midline as RNAi of *hid* or overexpression of the *Drosophila* inhibitor of caspases (Diap1), which blocks apoptosis (Figure 1D and Figure 1I-1L). The midline increase with *rst* RNAi was lower than with *hbs*, but still highly significant (Figure 1I-1L). To further study the role of Hbs and Rst, we quantified cell elimination events using a live imaging set-up for the pupal notum, consisting of a glass window inserted into the pupal case, which allows recording of cell dynamics over 700 minutes across developmental timepoints from 18 to 30 hours after pupa formation (APF) (Figure 1A).

The presence of pannier-driven nuclear RFP and presence of membrane GFP (*E-cadherin::GFP*) facilitated the tracking of cell elimination (dying cells, yellow spots) in combination with RNAi (Figure 1M). Knock-down of *hbs* or *rst* in the *pannier* domain reduced cell delamination within the midline region compared to the control (RNAi of *white*) (Figure 1M-1N and Video 1 - Video 2). The effect was less strong compared to RNAi of *hid*, which almost completely blocked cell elimination during the imaged time window. This result suggests that in addition to reduced cell delamination, other factors contribute to the increased midline observed in adult *hbs* RNAi flies, which is comparable to *hid* RNAi (Figure 1D and Figure 1I).

Interestingly, knock-down of *hbs* and *rst* in the notum led to increased cell elimination at the peripheral region suggesting that reduced Hbs levels may lead to altered tissue dynamics (Figure 1M-1O). Together, the results showed that knock-down of Hbs and Rst led to a strongly altered pattern of cell selection in the developing epithelium, characterized by reduced cell removal at the midline and elevated cell death in peripheral areas. The observed cell delamination at the periphery could arise from altered mechanical forces across the notum. The findings demonstrate that the cell adhesion molecules Hbs and Rst are required for epithelial remodelling and correct formation of adult dorsal thorax structures, potentially by shaping mechanical forces or cell intercalation dynamics.

### Hbs and Rst shape global and local epithelial remodelling

Rst and Hbs have been shown to accumulate at the interface between hair cells and inter-hair cells during fly wing margin cell development, cells favoured to survive several apoptotic waves (Takemura and Adachi-Yamada, 2011). We therefore hypothesized that Rst/Hbs expression could be predictive of areas of cell survival versus cell elimination in the notum, potentially by reducing stress or facilitating signalling in certain areas.

To test this, we set out to record Hbs and Rst localization in the developing notum in real time, taking advantage of specific knock-in reporter lines that allow monitoring of dynamic Hbs and Rst expression as EGFP-tagged fusion proteins (Nagarkar-Jaiswal *et al*., 2015). Before, we confirmed the correct Hbs::GFP and Rst::GFP localization in wing imaginal discs along the dorsal-ventral (DV) boundary and in the notum region of the wing disc (Dworak *et al*., 2001; Linneweber, Winking and Fischbach, 2015) (Supplementary Figure 3A-3D). Interestingly, Hbs and Rst seem to localize at cell adherence junctions in regions predicted to give rise to sensory organ precursors (Supplementary Figure 3E-3F).

We then started notum recordings at 18-20 APF and noted that both Hbs::EGFP and Rst::EGFP were strikingly absent from the forming midline and enriched in cells located outside the midline region (Figures 2A-2H). This pattern was highly consistent and quantification of signal across the notum yielded a clear minimum of Hbs and Rst signal in the area of cell elimination (midline) (Figures 2C and 2G). However, their distribution outside the midline was not identical: Hbs was strongly expressed in bristle cell sockets (BcSs) with decreasing levels the farther cells were located from the central BcS (Figure 2B and Figure 2D). Rst expression was more uniform in peripheral areas but appeared often selectively enriched in certain cell junctions compared to others (Figures 2F and Figure 2H). In order to unequivocally correlate Hbs levels to one cell and track its fate in areas of differential cell compaction, we next imaged Hbs dynamics in *hbs-Gal4* flies driving nuclear GFP (*UAS.nlsGFP*). As previously observed with the reporter lines, *Hbs-Gal4* driven GFP appeared low in midline cells from 18-22h APF, only reaching equal expression to peripheral cells by 30h APF (Figures 2 I-K). We could not perform similar experiments for Rst due to the unavailability of suitable *rst-Gal4 lines*.

**Figure 2.**
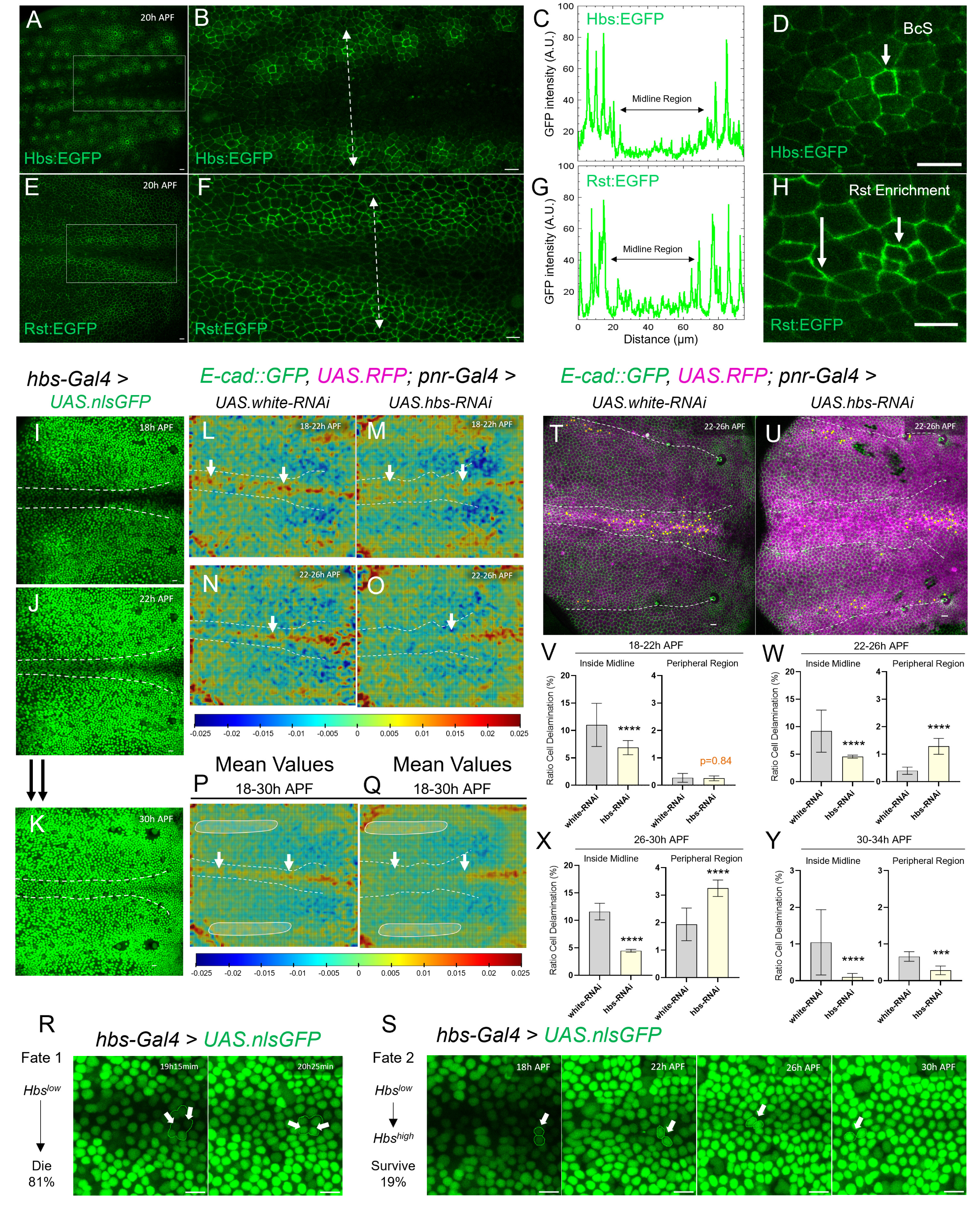
Hbs and Rst drive global and local epithelial remodelling. (A–H) Z-projections of live pupal nota at 20h APF expressing a Hbs reporter (A) or a Rst reporter (E). Close-up views of the midline region (outlined by white rectangles in A and E) are shown for Hbs-EGFP (B) and Rst-EGFP (F). Intensity profiles of Hbs (C) and Rst (G) measured along the white dashed line crossing the midline region. Close-up views showing Hbs localization (D), and Rst enrichment (H). Scale bars: 10 µm. (I–K) Z-projections of live pupal nota at 18h (I), 22h (J), and 30h (K) APF from 700min movies expressing GFP (green) in the Hbs domain. The midline region is highlighted in white. Scale bars: 10 µm. (L–Q) Averaged PIV vector fields showing tissue convergence rates from live imaging movies of control (*white* RNAi) and *hbs* RNAi pupae over matched time intervals. For *white* RNAi: 18–22h APF (L), 22–26h APF (N), and average from 18–30h APF (P). For *hbs* RNAi: 18–22h APF (M), 22–26h APF (O), and average from 18–30h APF (Q). Red regions indicate high convergence; blue regions indicate low convergence. White arrows highlight reduced cell convergence in the midline region upon *hbs* RNAi compared to control. Circles highlight increased convergence in the periphery upon *hbs* RNAi. (R–S) Close-up views of two distinct cell fates in the pupal notum midline region. Fate 1 (R) – Low Hbs-expressing cells elimination (arrows indicate delaminating cells). Fate 2 (S) – Low Hbs-expressing cells (white arrows) dynamically upregulate Hbs over time. Scale bars: 10 µm. (T–U) Z-projections of live pupal nota at 22h APF from 700min movies used to quantify cell death from 22–26h APF in control flies (*white* RNAi) and upon *hbs* RNAi. Yellow cells indicate future dying cells within the pannier domain (RFP, magenta) and Ubi-Ecad::GFP (green). White dashed lines mark the midline. Scale bars: 10 µm. (V–Y) Quantification of cell delamination events (ratio) inside midline and in the periphery region across sequential developmental intervals. Time windows: 18–22h APF (T), 22– 26h APF (U), 26–30h APF (V), 30–34h APF (X). Bars represent standard error of the mean (SEM). Statistical analysis was performed using Fisher’s exact test against the control; ****p < 0.0001.

It is known that during the earlier stages of pupal notum formation, high cell compaction and cell elimination rates at the midline contrast with moderate cell compaction and low cell elimination at the peripheral region (Levayer, Dupont and Moreno, 2016; Moreno *et al*., 2019). In order to assess how Hbs expression affected tissue compression, we next employed particle image velocimetry (PIV) to track tissue dynamics in nota with suppressed Hbs expression (*hbs* RNAi) versus controls (*white* RNAi) (Figures 2L-2Q). The generated compaction rate maps (calculated as -divergence of the vector field) showed that when Hbs^high^ and Hbs^low^ territories were abolished due to RNAi, the midline compression zone failed to build up efficiently at 18-22h APF, compared to controls (Figures 2L-2M). Lower cell compaction at the anterior half of the midline also persisted at later time points (22-30h APF) (Figure 2N-2O). When we integrated the total tissue deformation over time, the plots confirmed the lack of midline compaction zone in *hbs* knock-down conditions and showed more extensive areas of moderate compression in peripheral regions compared to controls (Figure 2P and 2Q, circled areas). We found similar pattern of compaction upon *rst* RNAi (Supplementary Figure 3G-3I). This suggests that peripheral Hbs and Rst levels are needed to drive cellular flow towards the midline. The direct involvement of Hbs in notum remodelling was supported by the fact that Hbs^low^ expression in the center strongly correlated with high cell compaction, whereas Hbs^high^ levels in the periphery overlapped with areas of low or moderate compaction (Figure 2I and 2J vs Figure 2L and 2N).

Apart from the detected effect of Hbs expression on cellular flow across the notum, we next studied local effects at the midline. We observed that cells with very low Hbs expression at the midline were eliminated (81% of 50 tracked cells), especially when surrounded by cells with higher Hbs levels (Figure 2R, Video 3). Precisely 19% of Hbs^Low^ cells persisted and this fate was associated with gradual Hbs upregulation around 22h APF (Figure 2S, Video 4).

To better understand the link between mechanical forces, Hbs expression and cell elimination, we analysed the impact of Hbs knock-down on cell death in short time windows (Figure 2T and 2U). During the first 4h of pupal formation, suppressed Hbs expression caused a reduction in cell delamination rates at the midline (Figure 2V) without altering cell death in the periphery. From 22h on, the clear reduction of cell pruning at the midline continued, combined with ectopic cell extrusion in the periphery, potentially causing a “flow” to the peripheral region, as cells accumulate at the widening midline (Figure 2W) in *hbs* RNAi flies. In the later stages of notum development, *hbs* knock-down caused mainly reduced cell delamination at the midline, with increased peripheral cell death, still observable at 26-30h APF (Figure 2X). Finally, cell elimination comes to a halt, when Hbs expression turns uniform at the end of notum remodelling (30h APF) (Figure 2Y). Similar videos were recorded for *rst* knock-down, which showed similar patterns of cell death with increased peripheral cell elimination as observed with *hbs* RNAi (Supplementary Figure 3J-3M).

These findings suggest that high Hbs and Rst expression in the peripheral notum is necessary to establish a well-positioned central compression zone that likely acts as a driving force for mechano-sensitive cell elimination. At the midline, the absence of Hbs expression is highly predictive of cell elimination. The observation that higher Hbs level in midline cells favours their maintenance over low expressers bears resemblance to results obtained in the fly retina, where interommatidial cells with high Hbs/Rst interactions are kept, whereas interommatidial cells with reduced levels are being eliminated (Bao and Cagan, 2005). Apart from a local protective effect at the midline, we find that Hbs and Rst expression patterns may dictate the pattern of cell death across the notum by affecting cell cohesion and compaction rate.

Our results support that Hbs and Rst function both at the tissue scale by modulating cellular flow and locally at the midline to confer selective protection of retained cells, although the two effects may be linked to some extent.

Since abrogating Hbs/Rst territories by RNAi resulted in midline width defects, we next asked if homogenous Hbs overexpression would interfere with midline formation. Indeed, quantification of the adult notum midline in flies with *pannier-Gal4-*dependent or *apterous-Gal*4-dependent Hbs overexpression (*UAS.hbs*) resulted in an increased midline, similar to overexpression of Diap1, which blocks apoptosis (Figures 3A-3G). Live imaging of the pupal notum showed that Hbs overexpression reduced the ratio of cell elimination at the midline compared to controls (*UAS.lacZ*), while cell death at the periphery remained unchanged (Figures 3H-3K).

**Figure 3.**
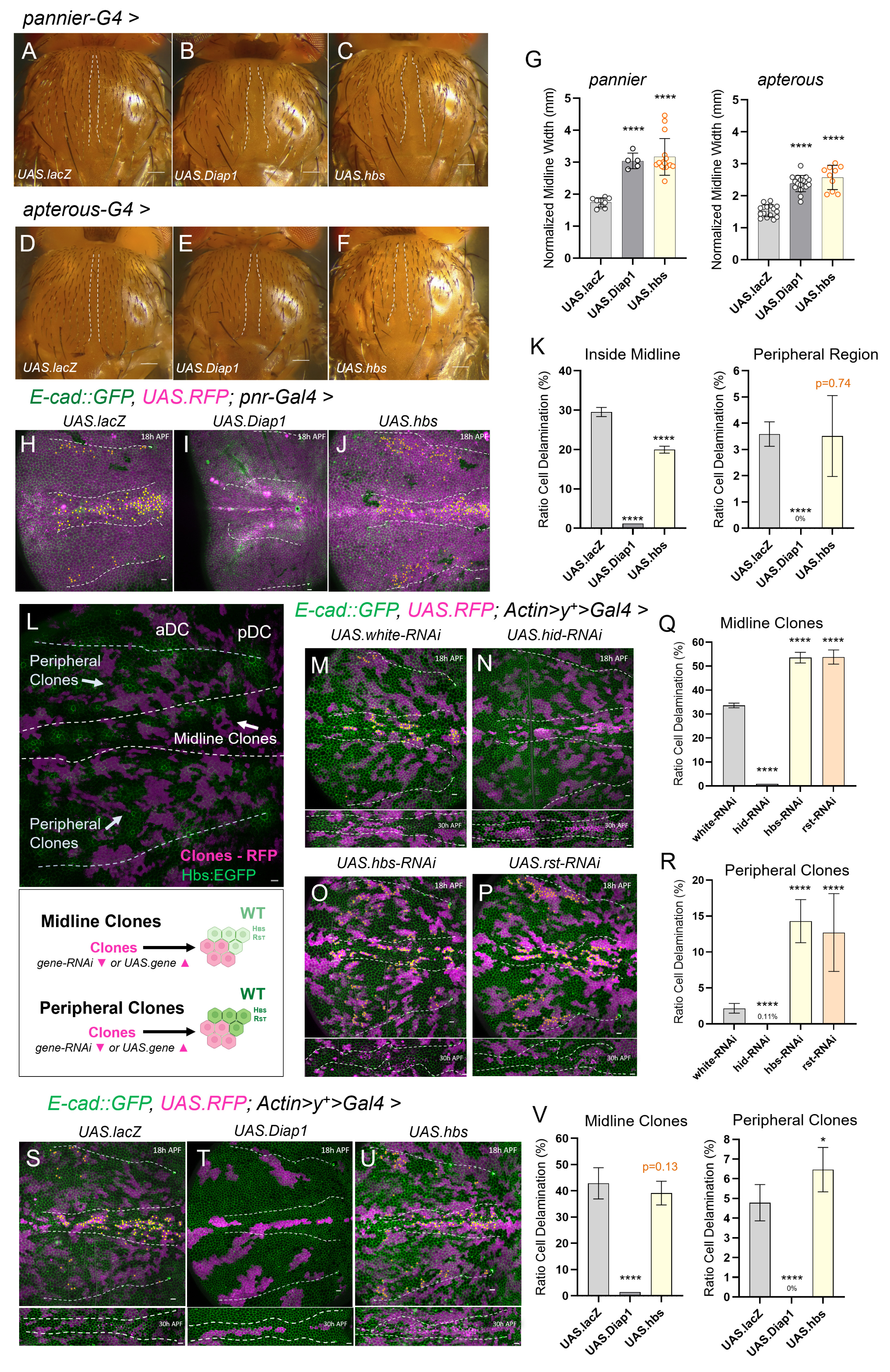
Hbs and Rst instruct cell elimination to fine-tune cell selection in the notum. (A–F) Adult notum images displaying midline phenotypes of control (*UAS.lacZ*), *UAS.Diap1* and *UAS.hbs* by *pannier* and *apterous Gal-4*. Scale bars: 10 µm. (G) Quantification of normalized midline width (mm) upon gene overexpression using *pannier-Gal4* (left) and *apterous-Gal4* (right). Each dot represents one adult fly thorax. Statistical significance was assessed by unpaired t-test or Mann-Whitney against the control (****p < 10⁻⁴). Bars represent standard deviation (SD). (H–J) Z-projections of live pupal nota at 18h APF (700min movies) with quantified cell death in control flies (*UAS.lacZ*) and upon *Diap1* and *hbs* overexpression. Yellow cells indicate future dying cells within the pannier domain (RFP, magenta). Cell membranes are labelled with *ubi-Ecad::GFP* (green). The midline region is outlined with white dashed lines. Peripheral regions analysed are delimited by the aDC and pDC bristles (white dashed lines). Scale bars: 10 µm. (K) Quantification of cell delamination events (ratio) inside the midline region (left) and in the periphery (right) over 12 hours (700 min; 18–30h APF). Bars represent the standard error of the mean (SEM). Sample sizes: *UAS.lacZ*: n = 3 nota, 1,167 cells (midline), 3,802 cells (outside); *UAS.Diap1*: n = 1 notum, 1,035 cells (midline), 1,784 cells (outside); *UAS.hbs*: n = 2 nota, 829 cells (midline), 2,836 cells (outside). Statistical analysis was performed using Fisher’s exact test against the control; ****p < 10⁻⁴. (L) Schematic illustrating two distinct cell interaction scenarios following clone induction in the pupal notum, based on the endogenous expression pattern of Hbs. Peripheral clones downregulating *hbs* interact with wild-type (WT) cells expressing high Hbs levels, while midline clones face WT cells with low Hbs. Conversely, *hbs*-overexpressing clones compete against high-Hbs WT cells in the periphery and low-Hbs WT cells in the midline. (M–P) Z-projections of live pupal nota at 18h APF (700min movies) with quantified cell death in control flies (*white* RNAi) and upon *hid* RNAi, *hbs* RNAi and *rst* RNAi. Yellow cells mark future dying cells within clones (RFP, magenta). Cell membranes are labelled with *ubi-Ecad::GFP* (green). Midline regions are outlined by white dashed lines at 18h APF and again at 30h APF below each image. Peripheral regions analysed are delimited by the aDC and pDC bristles (white dashed lines). Scale bars: 10 µm. (Q–R) Quantification of cell delamination events (ratio) in midline clones and peripheral clones over 12 hours (700 min; 18–30h APF). Bars represent SEM. Sample sizes: *white* RNAi: n = 3 nota, 469 cells (midline), 1,989 cells (outside); *hid* RNAi: n = 1 notum, 246 cells (midline), 942 cells (outside); *hbs* RNAi: n = 4 nota, 503 cells (midline), 2,385 cells (outside); *rst* RNAi: n = 2 nota, 289 cells (midline), 1,277 cells (outside). Statistical significance was assessed using Fisher’s exact test against the control; ****p < 10⁻⁴. (S–U) Z-projections of live pupal nota at 18h APF (700min movies) with quantified cell death in control flies (*UAS.lacZ*) and upon *Diap1* and *hbs* overexpression. Yellow cells indicate future dying cells within clones (RFP, magenta). Cell membranes are labelled with *ubi-Ecad::GFP* (green). White dashed lines mark the midline region at 18h and 30h APF. Peripheral regions analysed are delimited by the aDC and pDC bristles (white dashed lines). Scale bars: 10 µm. (V) Quantification of cell delamination events (ratio) in midline clones (top) and peripheral clones (bottom) over 12 hours (700 min; 18–30h APF). Bars represent SEM. Sample sizes: *UAS.lacZ*: n = 3 nota, 411 cells (midline), 2,024 cells (outside); *UAS.Diap1*: n = 1 notum, 70 cells (midline), 231 cells (outside); *UAS.hbs*: n = 4 nota, 481 cells (midline), 2,345 cells (outside). Statistical significance was assessed using Fisher’s exact test against the control; ****p < 10⁻⁴.

To understand if local alterations in the Hbs and Rst code would be sufficient to alter cell fate, we created RFP-marked *hbs* and *rst* RNAi clones surrounded by wild-type (wt) notum cells (RFP negative) using a heat-shock inducible flippase together with an *Actin>stop>Gal4* flip-out cassette. The random flip-out produced RNAi clones within the midline (midline clones) and in the periphery (peripheral clones) (Figure 3L). We then quantified cell delamination ratios in *hid* RNAi, *hbs* RNAi, and *rst* RNAi clones and control clones (*white* RNAi) under the *actin* promoter in the Hbs/Rst^Low^ midline or in the Hbs/Rst^high^ periphery (Figures 3M-3P; Video 5 – Video 6). In accordance with the previous fate tracking of *hbs* low cells, *hbs and rst* RNAi clones located at the midline showed significantly higher elimination rates compared to controls (*hbs* RNAi: +20.1%; *rst* RNAi; +18.82%) (Figure 3Q), confirming that low Hbs levels destined midline cells for elimination (Figure 2R). In comparison, *hid* RNAi clones showed no cell delamination as expected.

Interestingly, midlines with mosaic composition regarding Hbs or Rst all developed an irregular, meandering midline by 30h APF, which was not observed with control or *hid* RNAi (Figure 3M-3P, below each image) indicating that borders with strongly heterogenous Hbs/Rst levels may profoundly impact cell rearrangement or adhesion required to form the midline structure.

In the notum periphery, *hbs* and *rst* knock-down clones showed a 5-fold increased delamination rate (*hbs* RNAi: +12.18%; *rst* RNAi: +10.68%) (Figure 3R), in a zone where cell elimination is normally rare suggesting that Hbs/Rst protect against cell pruning.

Subsequently, we tested clonal overexpressing of Hbs, Diap1 or control β-Galactosidase (*UAS.lacZ*) in the midline versus the notum borders (Figure 3S-3V). Based on the fact that cells, which underwent Hbs upregulation could persist at the midline (fate 2, Figure 2S), we hypothesized that providing midline cells with an extra dose of Hbs may render them more resistant against cell delamination. However, we found that Hbs-overexpressing clones were not protected and showed similar elimination rates compared to controls, whereas Diap1-overexpression provided full protection (Figure 3V). Interestingly, Hbs overexpression in peripheral clones did cause a mild increase in delamination rates (+1.68%) (Figure 3V) indicating that the overexpression of Hbs likely produced considerably higher Hbs levels than the ones normally observed in the periphery. Indeed, previous findings had shown that Hbs overexpressing clones in the epithelial wing disc adopt a very indented morphology, displaying a higher cell mixing index (Tsuboi *et al*., 2017). It therefore appears that Hbs levels among neighbouring cells need to be highly in sync as any relative differences may affect cell cohesion properties.

Together, these findings reveal that the Hbs/Rst expression landscape in the notum is required to direct a correct cell elimination pattern. Hbs/Rst may initially exert an impact to direct cell flow and shape the central epithelial compression zone, while at a later stage, activation of endogenous Hbs/Rst levels, in a subset of midline cells, confers protection against cell delamination in the central compression area, concomitant with the completion of epithelial remodelling. This latter step may serve to fine-tune cell selection and retain cells with adequate shape and adhesion properties for optimal organ size.

### Differential regulation of Hbs and Rst by EGFR/ERK signalling

It has been previously shown that cell crowding at the midline induces downregulation of the EGFR/ERK pathway and upregulation of pro-apoptotic gene *hid*, leading to cell elimination (Moreno *et al*., 2019). EGFR is known to be enriched in adherens junctions in the notum (Moreno *et al*., 2019), similarly to what we detected for Hbs and Rst, suggesting potential interactions among these membrane proteins (Figure 4A). Since Rst and Hbs have been previously found to regulate EGFR signalling during *Drosophila* germline stem cell differentiation (Ben-Zvi and Volk, 2019), we sought to investigate if Hbs/Rst regulated cell elimination in parallel or upstream of EGFR/ERK signalling (Figure 4A). To address this question, we utilized a live sensor of ERK activity (*miniCic::mScarlet*) (Valon *et al*., 2021), which shifts from nuclear to cytoplasmic localization upon ERK phosphorylation (Moreno *et al*., 2019). Accordingly, nuclear mScarlet (magenta) is detected in cells that have low ERK activity, whereas the appearance of magenta signal in the cytoplasm indicates high ERK activation (Figure 4B). We detected ERK activity along the DV boundary of wing imaginal discs and at predicted SoP sites, as previously shown (Moreno *et al*., 2019), which correlated with sites of high Hbs and Rst expression (Supplementary Figure 4A-4D and Supplementary Figure 3A-3D). In the pupal notum, low ERK activation at the midline and in small patches in the periphery, overlaps with Hbs^low^ and Rst^low^ signal, respectively at 18APF (Figures 4C and 4D, arrows and close-up view). On the other hand, most peripheral areas with high ERK activity also showed high Hbs and Rst levels (arrowheads). This pattern was very consistent, although we occasionally found a few cells where this correlation was interrupted (circled cells). By the end of notum development, the entire notum displayed high levels of ERK and widespread Hbs and Rst expression (Supplementary Figure 4E-4F).

**Figure 4.**
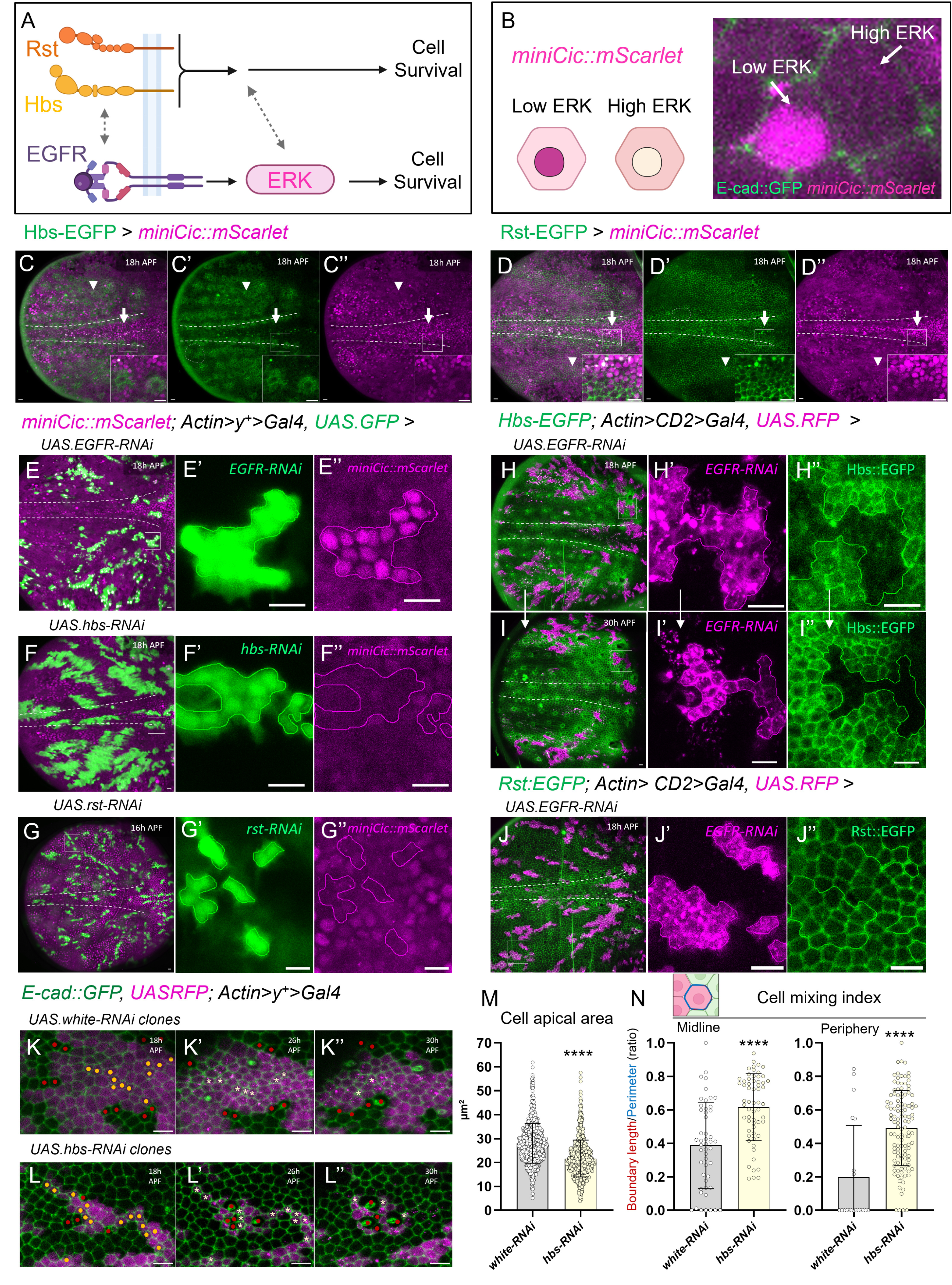
Hbs and Rst are Differentially Regulated by EGFR/ERK Signalling. A) Schematic model summarising the proposed roles and potential interactions between Hibris (Hbs), Roughest (Rst), and the Egfr/Erk signalling pathway during notum epithelium remodelling. B) Schematic illustrating miniCic::mScarlet reporter activity (magenta), which inversely correlates with Erk signalling. High miniCic signal indicates low Erk activity; low miniCic signal reflects high Erk activation. C–D) Z-projections of live pupal nota at 18h APF expressing either Hbs::EGFP and Rst::EGFP reporters (green), together with the miniCic::mScarlet reporter (magenta, inversely correlated with Erk activity). Close-up views highlight the correlation between Hbs, Rst and ERK activity inside and outside the midline region. White arrows highlight regions of low Hbs/Rst levels and low Erk activity and arrow heads regions of high Hbs/Rst levels and high Erk activity. Grey dashed circles indicate rare regions where Hbs and Rst expression levels appear to be inversely correlated with Erk activity. Scale bars: 10 µm. E–G) Z-projections of live pupal nota at 16/18h APF showing Erk activity (miniCic::mScarlet, magenta) in clones (GFP, green) upon Egfr RNAi, hbs RNAi, or rst RNAi. Close-up views of clone-containing regions (white rectangles) are shown to the right for each panel. Scale bars: 10 µm. H–I) Z-projections of live pupal nota expressing the Hbs::EGFP reporter in the presence of Egfr knockdown clones (magenta) at 18h and 30h APF. Close-up views of the same clone-containing regions (white rectangles) are shown on the right. Scale bars: 10 µm. J) Z-projection of live pupal notum at 18h APF expressing Rst::EGFP (green) and Egfr knockdown clones (magenta). Close-up view of a clone-containing region (white rectangle) is shown to the right. Scale bars: 10 µm. K–L) Close-ups of control peripheral clones (*white* RNAi) and *hbs* RNAi peripheral clones from live pupal notum movies (Figure 3M and 3P). Images show the same clones over time at 18h (K, L), 22h (K’, L’), 26h (K’’, L’’), and 30h APF (K’’’, L’’’). Yellow clone cells at 18h indicate future dying cells; red cells mark neighbouring WT cells. Asterisks indicate clone cells moments before elimination. Scale bars: 10 µm. M-N) Quantification of cell apical area (µm²) and cell mixing index (defined as the fraction of a cell’s perimeter is in contact with non-clonal wild-type neighbours across the clone boundary and calculated as boundary length (red on scheme) / perimeter (blue on scheme) in control clones (*white* RNAi) and in *hbs* RNAi clones inside the midline region and in the periphery at 18h APF. Each dot represents one clone cell. Sample sizes: Cell apical area of all clone cells; *white* RNAi, n = 912 cells; *hbs* RNAi, n = 746 cells; Cell mixing index of future dying cells; *white* RNAi, n = 49 cells midline and n = 20 cells periphery; *hbs* RNAi, n = 60 cells midline, n = 116 cells periphery. Statistical significance was determined using the Mann–Whitney test; ****p < 0.0001.

To determine whether Hbs and Rst regulate EGFR/ERK signalling we assessed ERK activity in *hbs* or *rst* RNAi clones. *hbs* and *rst* knock-down cells still displayed high ERK activity, in contrast to *Egfr* RNAi clones that cannot activate ERK (Figures 4E-4G), indicating that Hbs and Rst do not function upstream of EGFR/ERK. To understand if the EGFR/ERK axis in turn may regulate Hbs or Rst, we assessed Hbs/Rst levels in EGFR knock-down clones. Interestingly, we found that suppression of EGFR caused an almost complete loss of Hbs membrane signal in most cells analysed at 18h APF and 30h APF (Figures 4H-4I). As an exception, few peripheral clones spanning bristle cells retained Hbs expression, suggesting an alternative regulation of Hbs in bristle formation (Supplementary Figure 4G-4H). In stark contrast to Hbs, Rst expression was not reduced by *Egfr* RNAi (Figures 4J), showing that the EGFR/ERK pathway regulates Hbs but not Rst expression.

The pro-survival EGFR/ERK signalling is required to prevent the activation of the pro-apoptotic *hid* that promotes caspase activation and cell death (Moreno *et al*., 2019). Given that EGFR/ERK signals are also required to maintain robust Hbs expression, it is envisageable that lower Hbs levels prepare cells for being compressed and extruded, by reducing adhesion and membrane resistance to compression. In support of this, we found that retrieved clones with compromised Hbs levels were generally smaller and showed a convoluted outline (Figure 4K-4L). As such, cells in *hbs* RNAi clones presented a consistently smaller apical surface when imaged compared to control clones (*white* RNAi) (Figure 4M). They also showed increased mixing/intercalation with wild-type cells (Figure 4L, red dots) as found in the wing disc (Tsuboi et al. 2017) and elevated cell delamination (yellow dots). Indeed, *hbs* RNAi cell clones showed an higher cell mixing index in comparison to control clones (Figures 4N). These findings suggest that high Hbs levels contribute to mechanical resistance whereas dynamic downregulation of Hbs in midline cells, potentially due to decreasing EGFR/ERK activity, may facilitate cell shrinkage and detachment.

Recent findings have revealed a role for cell geometry independent of mechanical forces (Cachoux *et al*., 2023), where smaller cells compared to their neighbours were preferentially eliminated in a Notch-dependent manner. As there is some evidence that E-cadherin can regulate Notch signalling in a way to direct Rst to specific IPC borders (Grzeschik and Knust, 2005), it is possible that the Hbs interaction partner Rst could be regulated by Notch.

## Conclusion

Overall, our screen has pointed out new regulatory components that refine cell elimination in the developing notum. The gained insight on Hbs and Rst function in notum remodelling further enhances our understanding of how organs are shaped. In particular, Hbs emerges as a crucial regulator that guides cell dynamics at the tissue level and regulates final selection of midline cells exposed to mechanical stress. We propose that Hbs acts as an integrating platform to convey both mechanical signals from compaction-dependent competition via EGFR/ERK and likely other cell adhesion or shape signals, potentially via heterophilic interaction with Rst. Hbs and Rst function is evolutionary conserved, where immunoglobulin superfamily members such as nephrin (encoded by *hbs* human homolog NPSH1) and Neph1 (encoded by *rst* homolog KIRREL) form specialized cell–cell interactions essential for the formation of the three-dimensional epithelial glomerular podocyte architecture (Simons, Hartleben and Huber, 2009). Mutations in NPHS1 are associated with congenital nephrotic syndrome (CNF) in humans (Tryggvason, Patrakka and Wartiovaara, 2006). Mice lacking *Nphs1* or *Kirrel* exhibit severe proteinuria, fail to form the slit diaphragm, and die shortly after birth (Donoviel *et al*., 2001). The overexpression of NPSH1 has also been found to drive cell aggregation in cancer cells (Khoshnoodi *et al*., 2003) and mutations in *KIRREL1* induce proliferative advantage to several human cancer cell lines, in a mechanism dependent on cell density levels (Paul *et al*., 2022). These findings raise the possibility that Hbs/Rst patterns may shape cell behavior and survival not only during epithelial remodelling, but also as a consequence of epithelial renewal or in the course of cancer formation.

Insight into the molecular mechanisms linking mechanical forces to cell adhesion via Hbs and Rst may therefore provide new avenues to restrict cancer growth in future.

## Methods

### Fly Husbandry, Genetics, and Clone Induction

All experiments were performed using *Drosophila melanogaster* fly strains maintained with standard husbandry techniques on Vienna standard cornmeal-molasses-agar media. All experiments were performed using female flies maintained at 25°C under a 12h light/dark cycle unless otherwise indicated. The flies’ health and immune status were not assessed, and they were not subjected to any prior procedures. Experimental fly lines are listed in Table 1; additional lines used in the screen are shown in Supplementary Table 1.

**Table 1:**
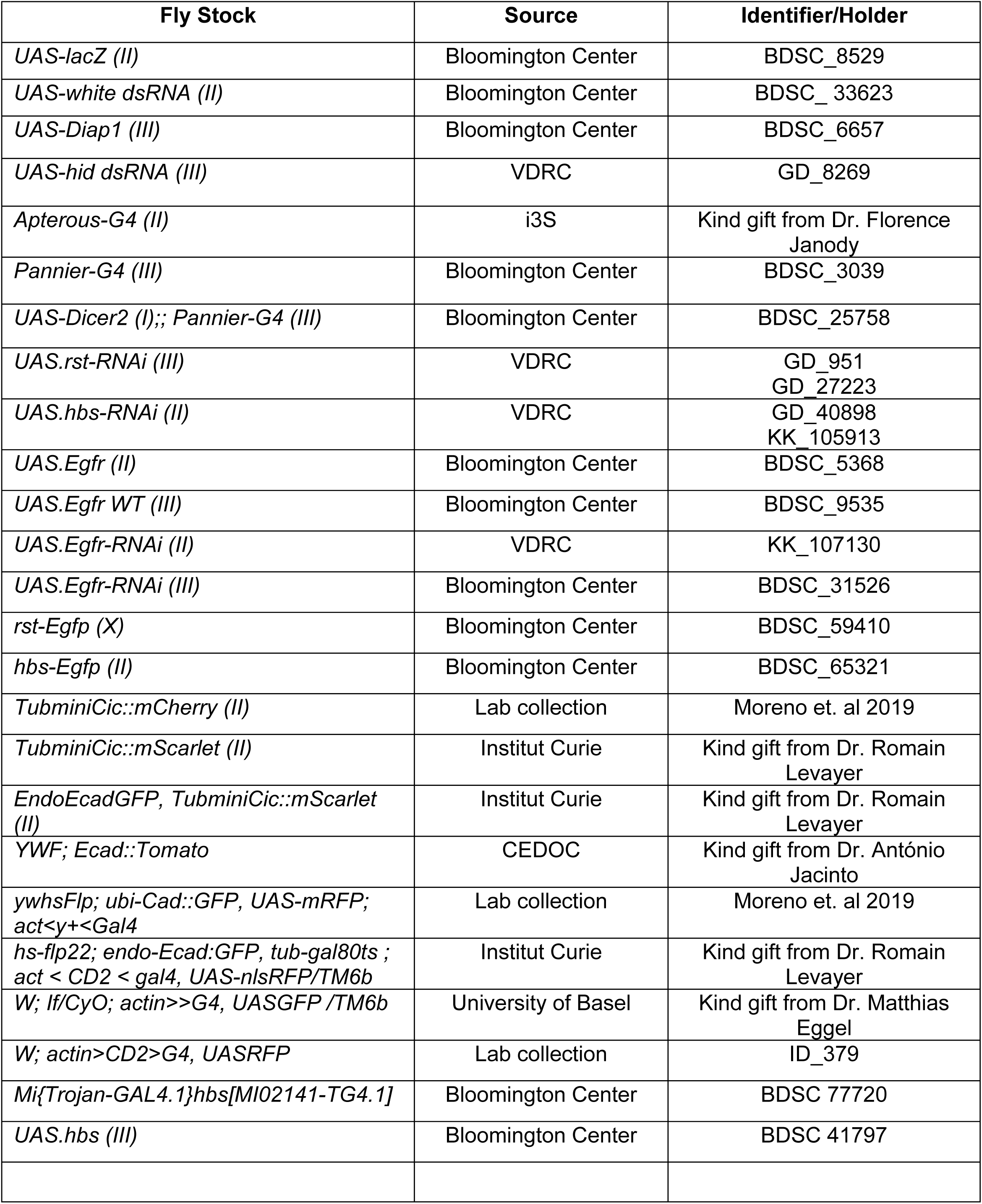
Fly stocks used in this study.

To identify regulators of the midline region formation, we performed an *in-silico* screen based on data from the Bristle Screen Database (Mummery-Widmer *et al*., 2009), a genome-wide RNAi screen in the Drosophila notum. Candidate genes were knocked down using UAS-RNAi lines and two notum-specific Gal4 drivers (*pannier-Gal4* and *apterous-Gal4*). Adult thoraxes were imaged using a Leica S9i camera with a 5.5xGreenough Stereozoom and 1.6x objective. he notum thorax images displayed are representative pictures for each studied phenotype of adult female flies raised at 25°C. Midline width was quantified as the mean of three measurements taken between the most central bristle rows at anterior, middle, and posterior regions, normalized to the distance between the two posterior supra-alar (pSA) bristles.

### Immunohistochemistry

Wing imaginal discs were dissected from wandering third instar larvae in PBS on ice, fixed in 4% formaldehyde (30 min), and permeabilized in 0.4% PBT. F-actin was labelled with Phalloidin (1:500, Molecular Probes); nuclei were stained with DAPI (1 μg/mL, Sigma). Samples were mounted in Vectashield (Vector Labs) and imaged with a Zeiss LSM880 using 20x (NA 0.8) dry or 40x (NA 1.4) oil objectives. Fluorescent genotypes are listed in Table 1.

### Live Imaging of the Pupal Notum and Clone Generation

To investigate the function of *hbs* and *rst*, RNAi clones were induced during early third instar using the FLP-out system in flies expressing either *E-cadherin::GFP* or *miniCic::mScarlet*. Clones were generated by applying a heat shock at 37°C for 15 min 48–72 hours prior to live imaging. White prepupae (0h after puparium formation, APF) were aged at 25 °C until 16–18h APF before dissection and imaging. For mounting, pupae were positioned between double-sided tape and coverslip spacers (assembled from four stacked #1 coverslips), dissected from the anterior end of the pupal case, and mounted in Halocarbon Oil 10S under a 20 × 40 mm #1.5 coverslip, sealed with nail polish.

### Image Acquisition and Cell Delamination Analysis

Live imaging was performed on a Zeiss LSM800 with fast Airyscan using a 40x oil objective (NA 1.4), acquiring Z-stacks (1 μm steps) every 5 min for 700 min (18–30h APF). Autofocus was maintained using *E-cadherin::GFP* plane as reference (Zen macro, Ellenberg Lab). Movies were captured near the scutellum, encompassing the midline and the aDC/pDC macrochaetae. Movies shown are maximum projections. Cell delamination was manually marked using the Cell Counter plugin in Fiji. Regions of interest (ROIs) were defined at movie onset based on the central bristle rows (midline) and the pDC bristle line (periphery) (Figure 1A). Only cells within each ROI during the entire recording were included in the analysis. Cell death probability was calculated as the ratio of extruded cells divided by the total initial cells per region (midline or periphery). Cells were considered “dying cells” if they or at least one of their daughter cells died before the end of the movie. Cell death quantification was not performed in regions posterior to the pDCs. Cell clones were manually segmented and identified in Fiji using cell contours defined by *E-cadherin::GFP* expression. The cell mixing index and apical cell area were quantified for control (*white* RNAi) and *hbs* knock-down clones at 18h APF. The cell mixing index was defined as the fraction of a cell’s perimeter that is shared with non-clonal (wild-type) neighbouring cells across the clone boundary and was calculated as the ratio between the boundary length shared with wild-type cells and the total perimeter of the clone (boundary length / perimeter). Cell apical area was measured in µm^2^ from cell contours in Fiji for both control and *hbs* knock-down clones.

### PIV Analysis

Tissue deformation was quantified using Particle Image Velocimetry (PIV) in MATLAB (PIVlab; has previously shown on (Levayer, Dupont and Moreno, 2016; Moreno *et al*., 2019). Two-pass analysis was performed with interrogation windows of 64 pixels (first pass) and 32 px (second pass), each with 50% overlap. Final vector field is composed of displacement vectors separated by 32 pixels (15.5 μm). Divergence was calculated on every 32px x 32px square and averaged over 240 or 700 min, as indicated in figure legends. Compaction rate (Convergence) is defined as ‘‘-Divergence’’ of the vector field (calculated on MATLAB). Averaged compaction rates were calculated by averaging on one PIV window the compaction rate over the full movie (700 min).

### Quantification and Statistical Analysis

Quantifications from adult thorax images are presented as mean ± standard deviation (SD). Normality was assessed using the Shapiro–Wilk test. Depending on the distribution, either parametric (unpaired t-test) or non-parametric (Mann–Whitney U) tests were applied. For live imaging-based cell death analysis, data are presented as mean ± standard error of the mean (SEM). To assess differences in binary cell fates (survival vs. death), statistical significance of cell delamination rates between conditions was determined using Fisher’s exact test, based on initial cell counts. Significance thresholds are indicated in the figure legends. All statistical analyses were performed using GraphPad Prism version 9.

## Acknowledgments

We thank the Bloomington Stock Center and the VDRC Stock Center for fly stocks; Romain Levayer, António Jacinto, Florence Janody, Matthias Eggel for sharing antibodies or fly stocks. We sincerely thank Catarina Brás Pereira for discussions, advice and support in several stages of this work. We thank Champalimaud Foundation’s fly facility and ABBE imaging platform for all the background work that contributed to the development of this project.

## Author Contributions statement

E.M. conceived and designed the project. M.F-P., M.A., C.R. and E.M. designed the experiments, M.F-P. conducted the experiments and analysed data; M.F-P and M.A. performed the multi-step screen data acquisition and analysis. M.A., C.R. and E.M. supervised the work. M.F-P and C.R. wrote the manuscript with contributions from the other authors.

## Supplementary Figure Legends

**Supplementary Figure 1.**
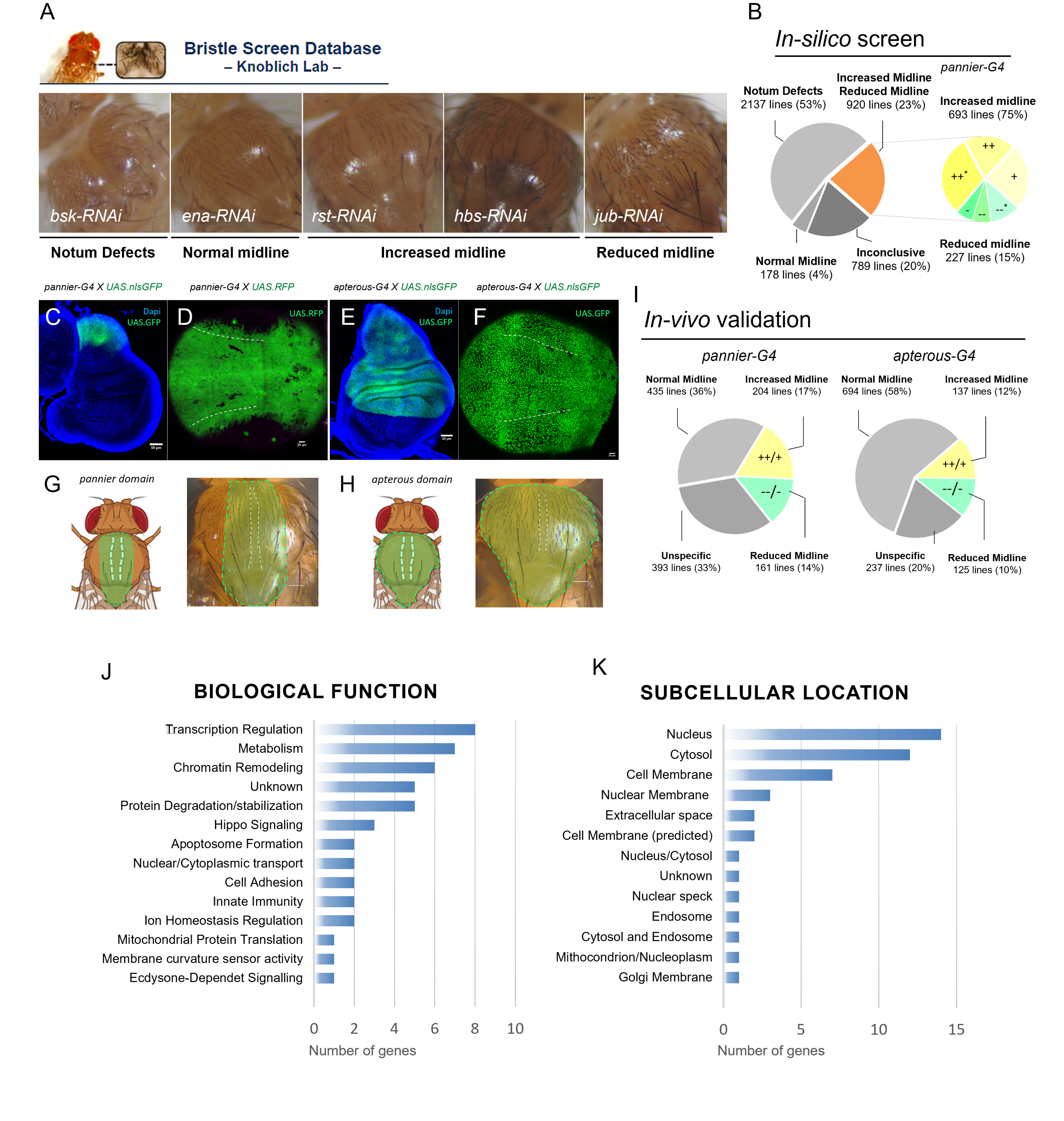
RNAi screen and characteristics of novel epithelial remodelling regulators. A) Examples of adult notum images analysed during the *in-silico* screen from the Bristle Screen Database. Each picture shows a typical example for each phenotypic category scored in our screen. Examples from left to right are *basket-RNAi* (Notum Defects), *enabled-RNAi* (Normal Midline), *roughest-RNAi* and *hibris-RNAi* (Increased Midline), *ajuba-RNAi* (Reduced Midline). The midline region is highlighted in white dashed lines. B) Recovered midline phenotypes in the “*in-silico* screen” based on RNAi activation with *pannier-Gal4*. Total number of RNAi lines and their relative frequency for each scored phenotype is displayed. RNAi lines scored as increased or reduced were further separated as: Increased strong (++) or intermediate (+) score, or Reduced strong (--) or intermediate (-) score. Asterisks mark RNAi lines that caused mild notum defects outside the midline. (C–H) Schematics illustrating the expression pattern of the two notum drivers used in the screen, *pannier* and *apterous*. The expression pattern in the wing disc is demonstrated for both drivers. Live pupa notum images showing *pannier* and *apterous* profiles at 18h APF. I) Recovered midline phenotypes in the validation screen step based on RNAi activation with *pannier and apterous-Gal4*. Total number of RNAi lines and their relative frequency for each scored phenotype and driver is displayed. RNAi lines scored as increased (++ or +) or reduced (-- or -). (J-K) Biological functions and subcellular locations of the novel 47 MDCD regulators. Gene characteristics were categorised according to their described function in the literature, utilising information available at FlyBase and UniProt websites.

**Supplementary Table 1:**
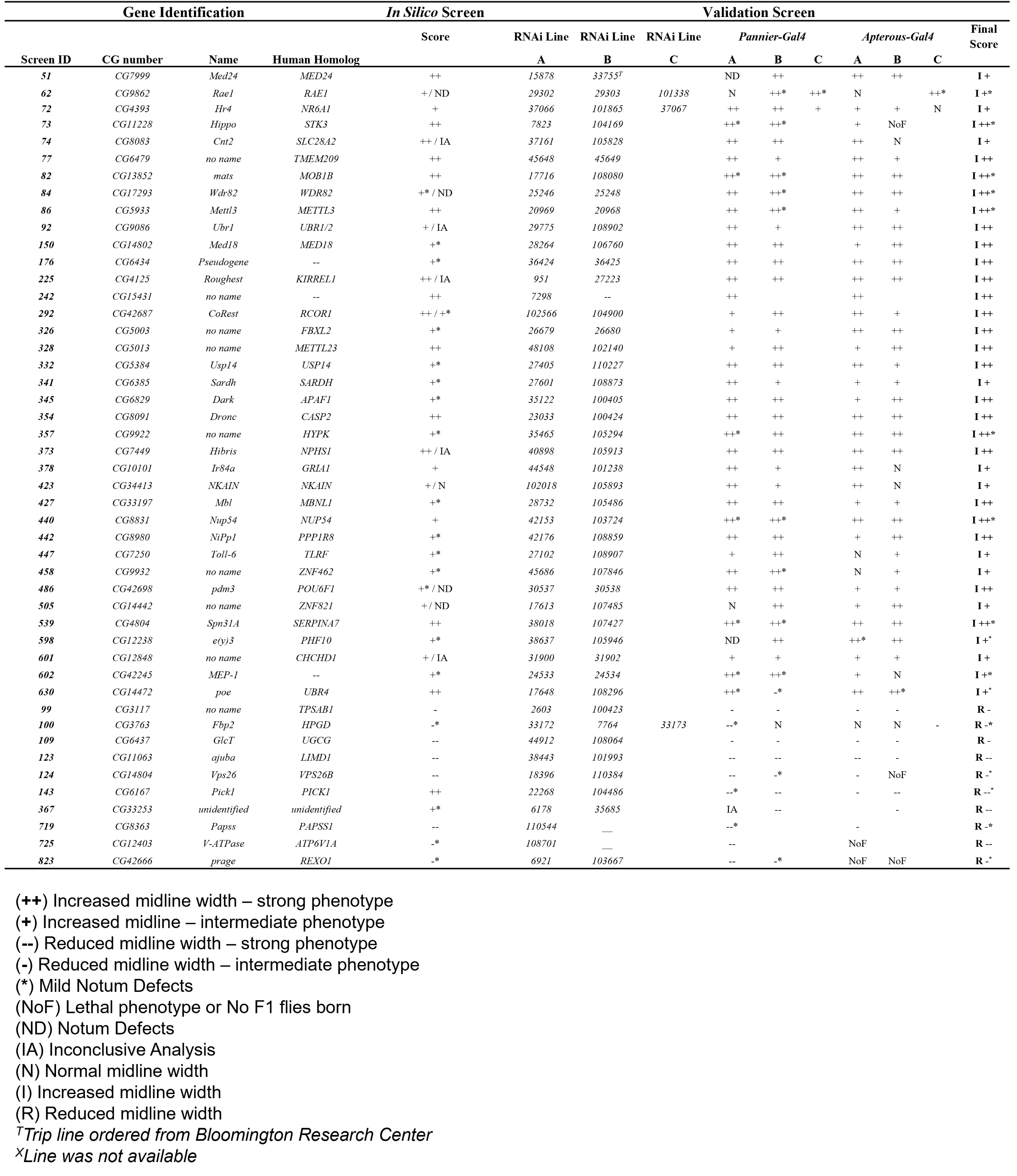
Screening result – Characteristics of novel epithelial remodelling regulators.

**Supplementary Figure 2.**
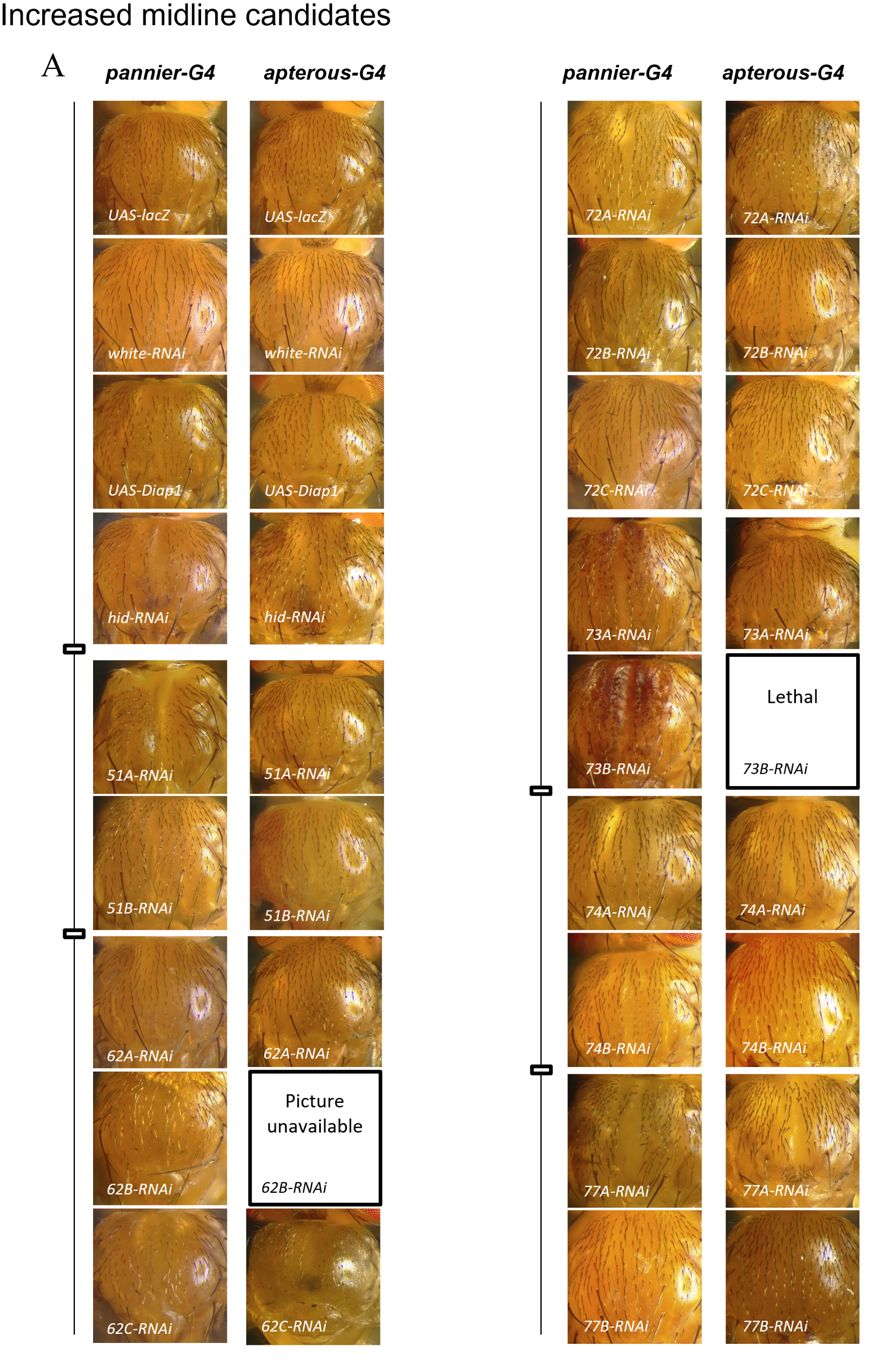
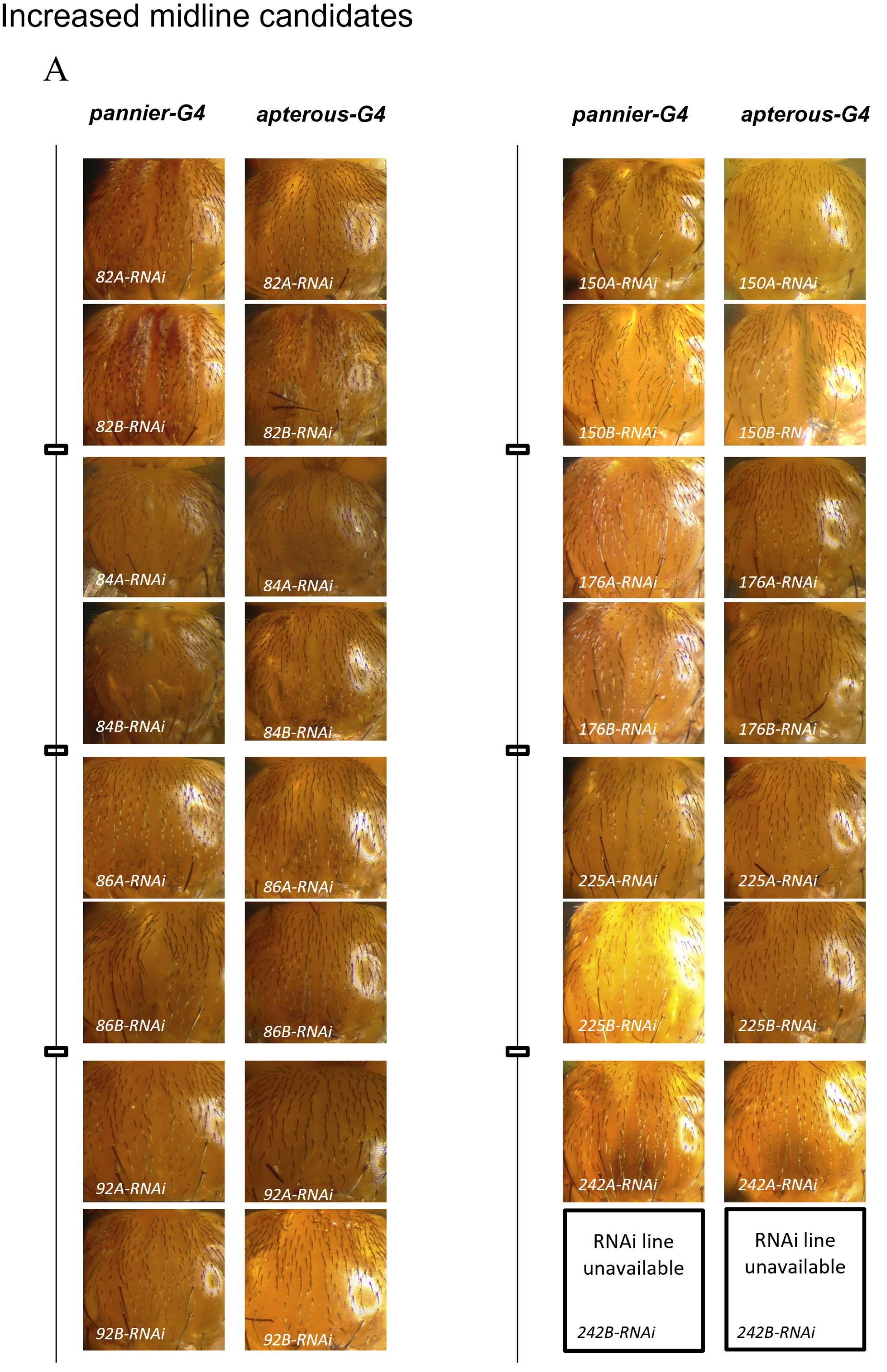
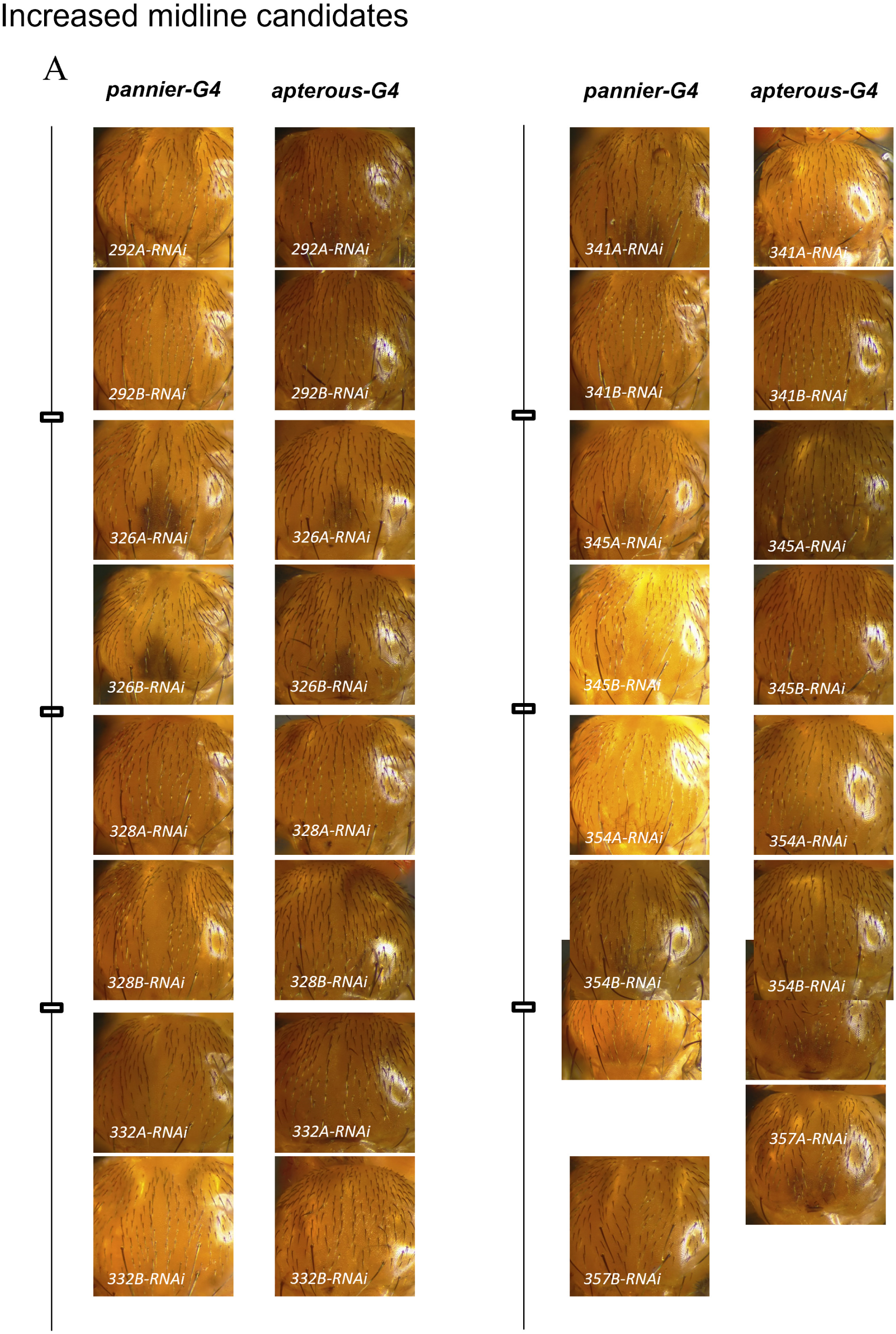
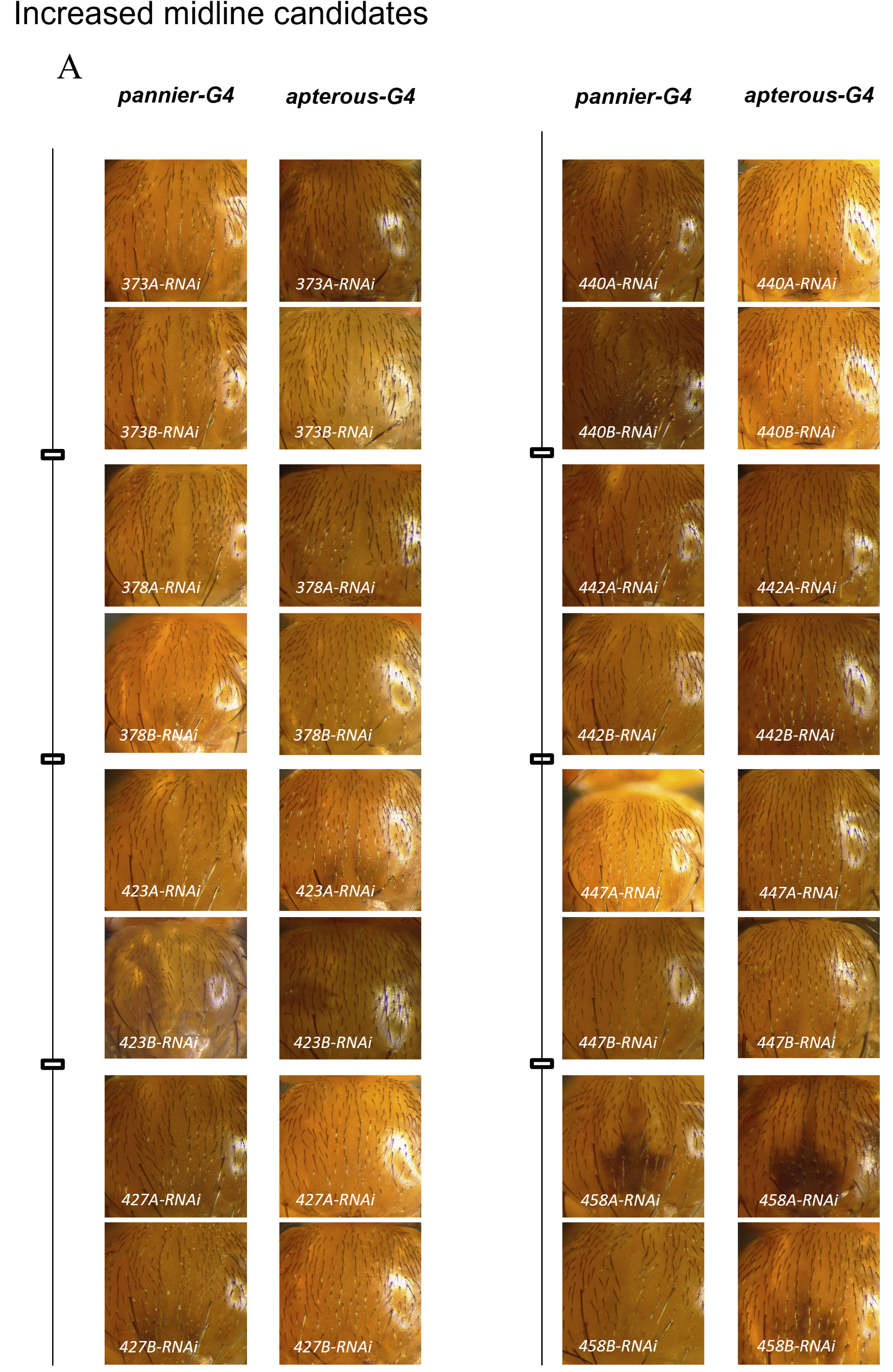
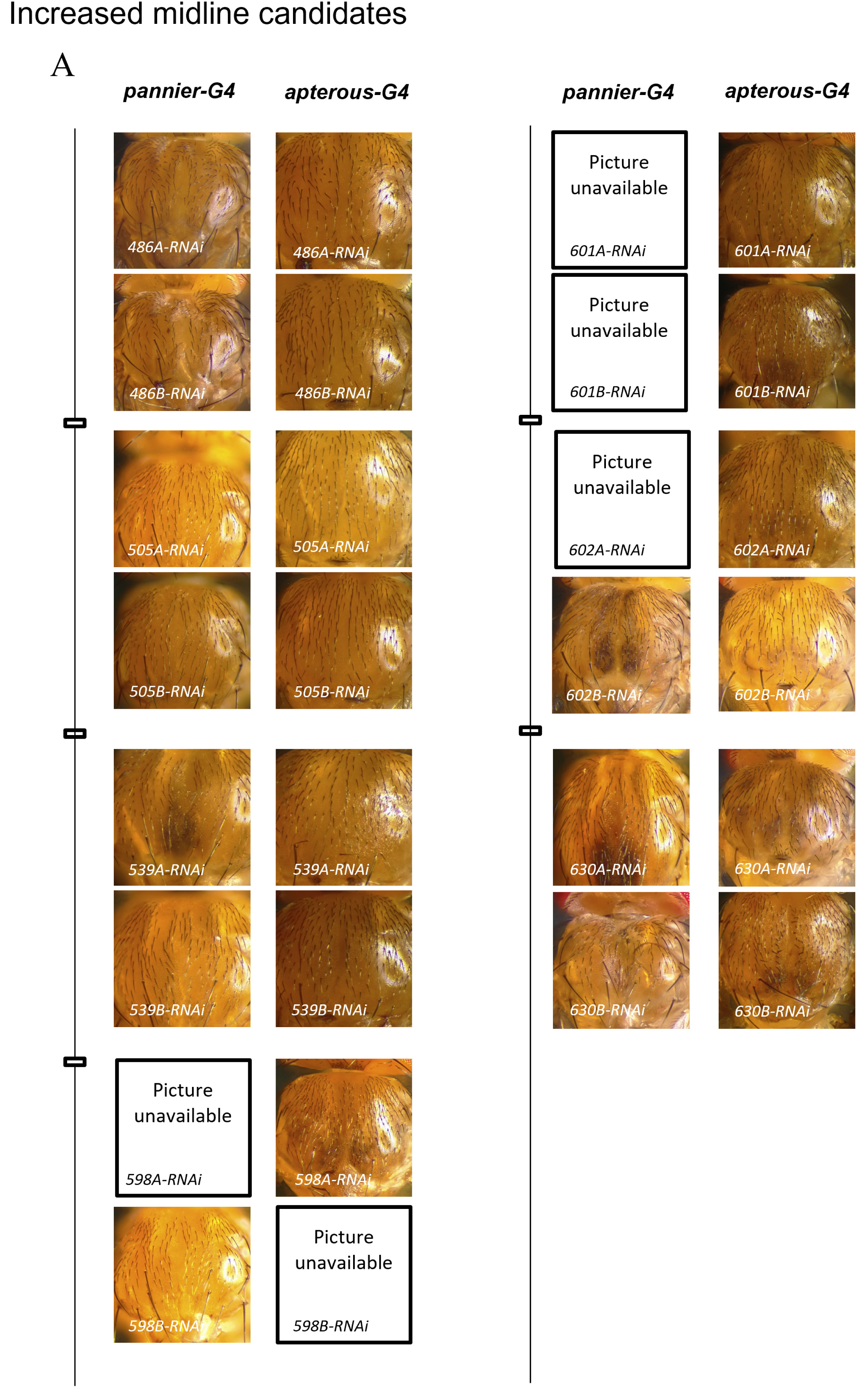
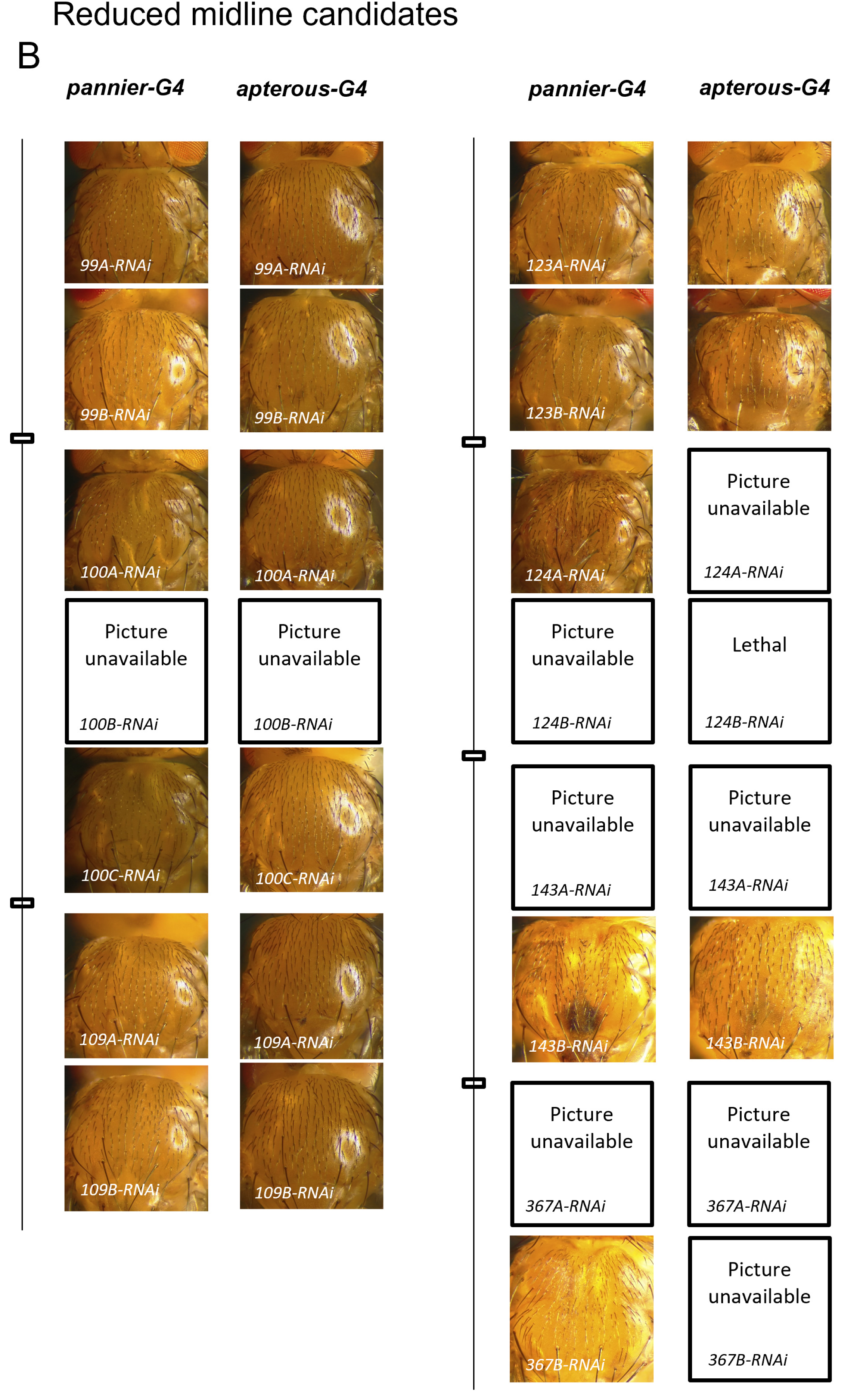
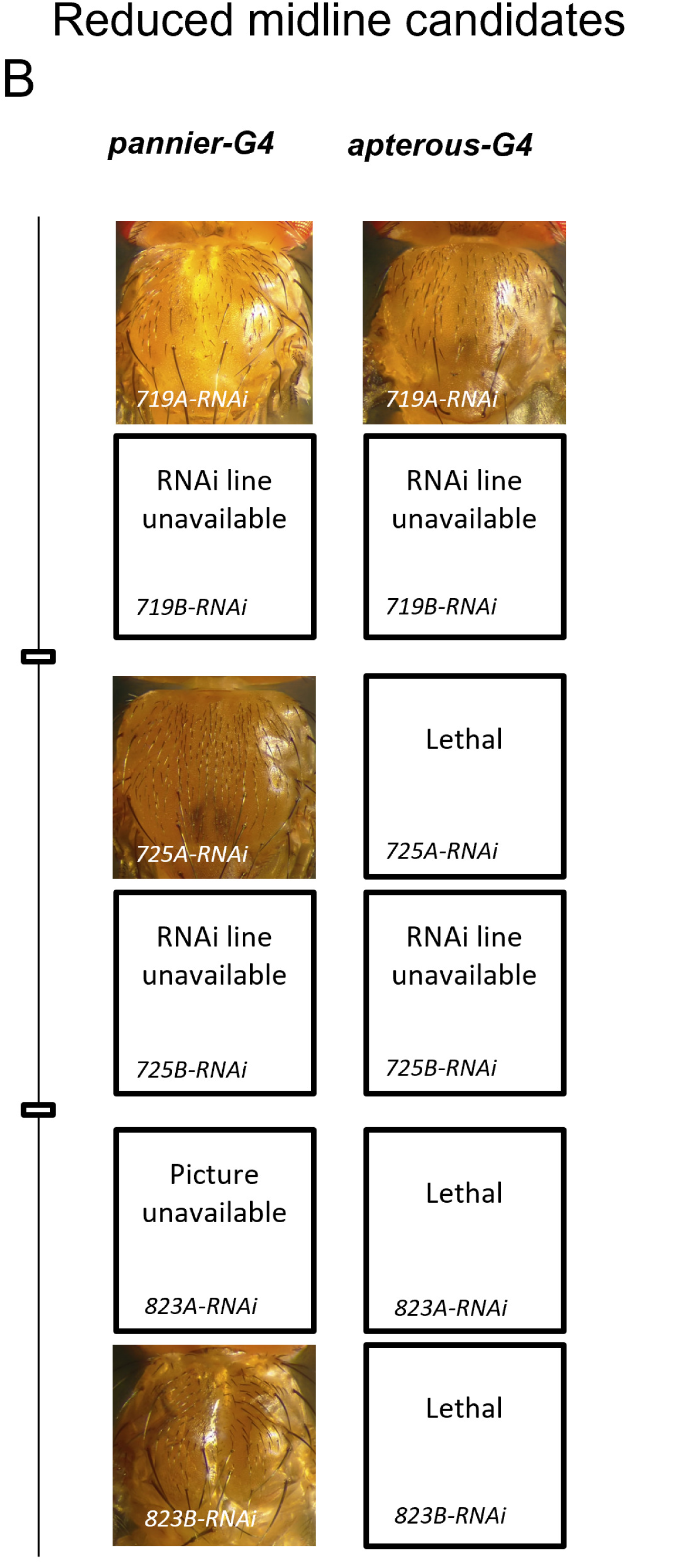
Notum phenotypes of the 47 validated candidates from the screen. (A-B) Snapshots of adult fly notum for the displayed genotypes. Representative pictures of all the new 47 candidates for each RNAi line tested and driver (*pannier* or *apterous)*, separated according to their screen ID (shown on Table 1). Increased midline width candidates(A). Reduced midline width candidates (B).

**Supplementary Figure 3.**
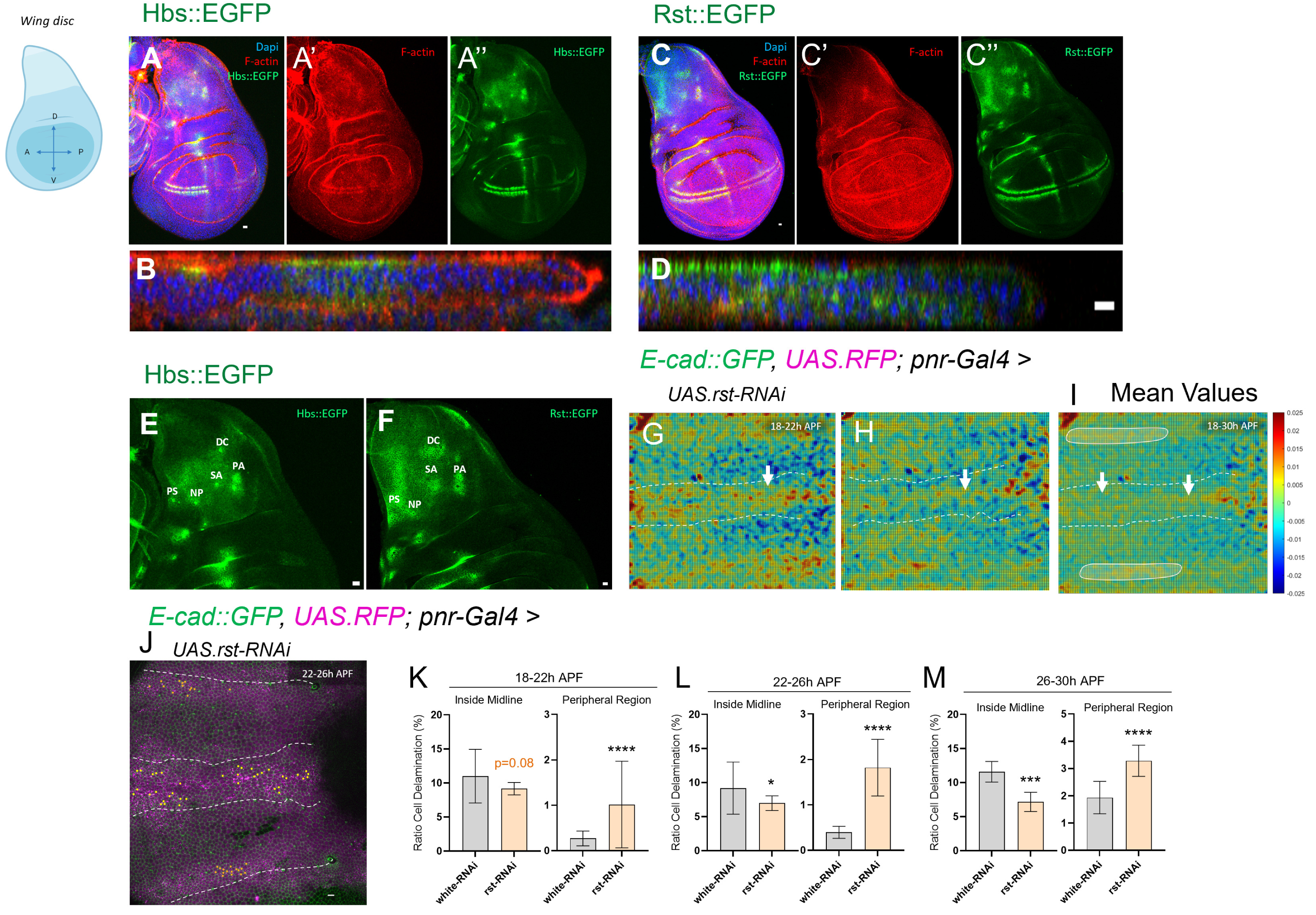
Hbs and Rst expression in the wing disc; Rst is necessary to drive global and local epithelial remodelling. A-D) Z-projections of wing imaginal disc at larvae L3 wandering stage expressing Hbs::GFP, Rst::GFP. Orthogonal views are shown below each reporter. Samples were stained for F-actin (phalloidin - red) and DAPI (blue). 4 tissues we used. Scheme illustrating wing disc orientation is shown on the right. Scale bars represent 10μm. (E-F) Close-up views of the notum region of the wing disc. Predicted BcSc containing regions are highlighted (DC, SA, PA, PS, NP). Scale bars: 10 µm. G-I) Averaged PIV vector fields showing tissue convergence rates from live imaging movies of *rst* RNAi pupae over matched time intervals. 18–22h APF (J), 22–26h APF (K) and average from 18–30h APF (L). Red regions indicate high convergence; blue regions indicate low convergence. Circles highlight extensive moderate convergence regions in the periphery upon *rst* RNAi compared to control (*white* RNAi from Figure 2L-2Q) (J) Z-projections of live pupal nota at 22h APF from 700min movies used to quantify cell death from 22–26h APF in control flies (*white* RNAi) and upon *rst* RNAi. Yellow cells indicate future dying cells within the pannier domain (RFP, magenta) and membranes *Ubi-Ecad::GFP* (green). White dashed lines mark the midline. Scale bars: 10 µm. (K–M) Quantification of cell delamination events (ratio) inside midline and in the periphery region across sequential developmental intervals. Time windows: 18–22h APF (N), 22– 26h APF (O) and 26–30h APF (P). Bars represent standard error of the mean (SEM). Statistical analysis was performed using Fisher’s exact test against the control; ****p < 0.0001.

**Supplementary Figure 4.**
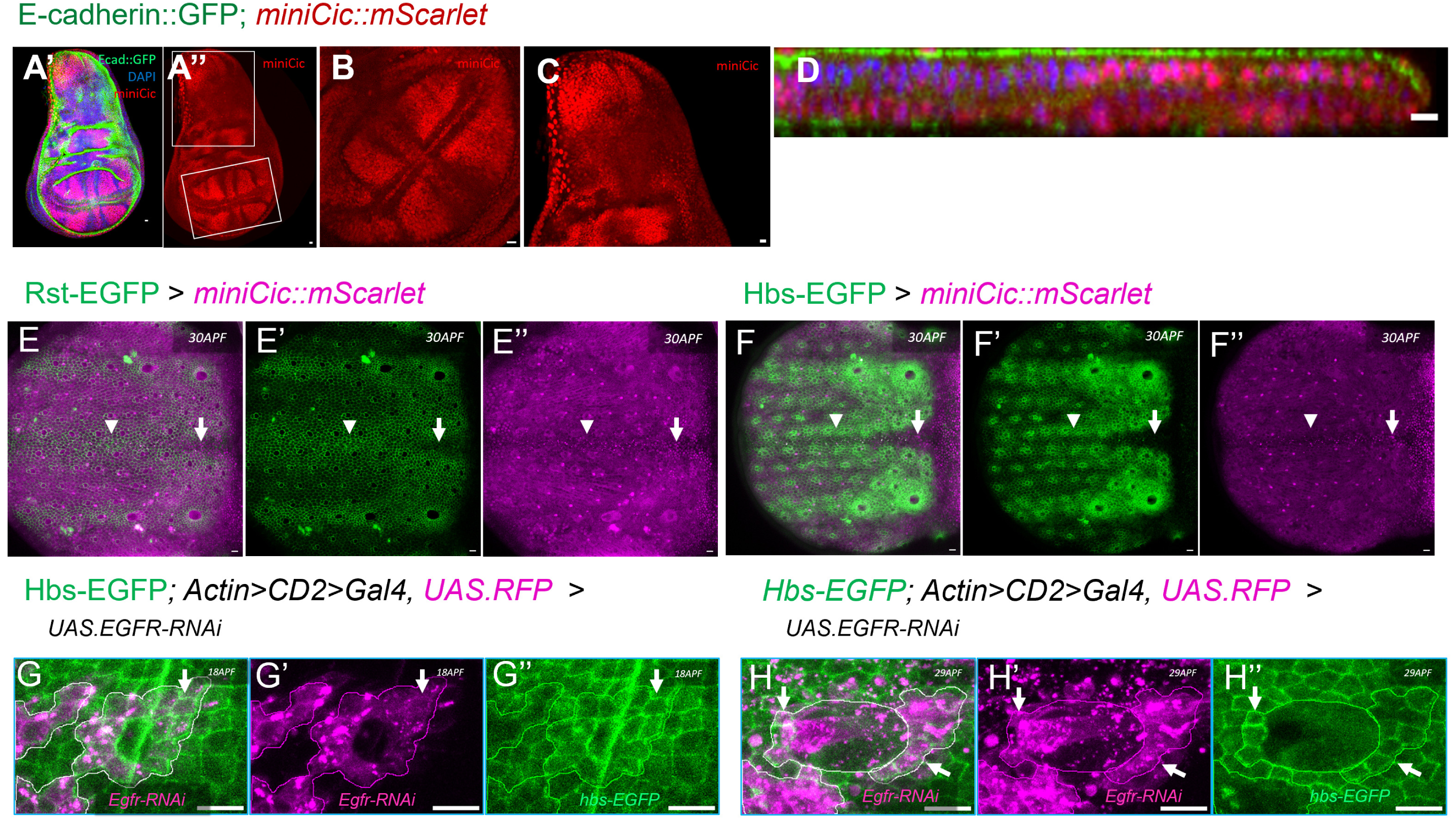
Hbs and Rst expression patterns largely correlate with EGFR/ERK activity in the wing disc; Rare exceptions of discrepancies between Hbs and ERK expression were observed near certain bristle cells in the pupa notum. (A-D) Z-projections of wing imaginal disc at larvae L3 wandering stage expressing *Ubi.Ecad::GFP* and ERK reporter (miniCic::mScarlet). Close-up views of the wing pouch (B) and notum (C) are shown. Orthogonal views of the notum region are shown on the right (D). Samples were stained for DAPI (blue). 10 tissues were used. Scale bars represent 10μm. E-F) Z-projections of live pupal nota at 30h APF expressing either Hbs::EGFP and Rst::EGFP reporters (green), together with the miniCic::mScarlet reporter (magenta, inversely correlated with Erk activity). White arrows highlight regions of low Hbs/Rst levels and low Erk activity and arrow heads high Hbs/Rst levels and high Erk activity. Scale bars: 10 µm. G-H) Z-projections of live pupal nota expressing the Hbs::EGFP reporter in the presence of EGFR knockdown clones (magenta) at 30h APF near bristle cells. Arrows point to clone-containing regions where EGFR does not affect Hbs signal or location. Scale bars: 10 µm.

## Video Legends

**Video 1 and Video 2. Hbs is necessary during notum epithelium remodelling, related to Figure 1**

Z-projections of live pupal nota at 18h APF (700min movies) with quantified cell death in control flies (*white* RNAi - video 1) and upon *hbs* RNAi (video 2). Yellow cells indicate future dying cells within the *pannier* domain (RFP, magenta). Cell membrane is marked with *ubi-Ecad::GFP* (green). White lines mark the midline region and delimit the peripheral region near the aDC and pDC bristles. Scale bars: 10 µm.

**Video 3. Low Hbs expressing cells are eliminated during notum epithelium remodelling, related to Figure 2**

Close-up view of low Hbs-expressing cells elimination inside the midline region during live pupal notum imaging. Arrows point to dying cells. Scale bars: 10 µm.

**Video 4. Low Hbs expressing cells can upregulate Hbs to survive during notum epithelium remodelling, related to Figure 2**

Close-up view of low Hbs-expressing cells inside the midline region gradually upregulating Hbs over time during live pupal notum imaging. Cells gradually upregulating Hbs are circled at 60-minute intervals to highlight progressive Hbs expression. Scale bar: 10 µm.

**Video 5 and Video 6. Hbs and Rst direct cell elimination to fine-tune cell selection during notum epithelium remodelling, related to Figure 3**

Z-projections of live pupal nota at 18h APF (700min movies) with quantified cell death in control flies (*white* RNAi – video 5) and upon *hbs* RNAi (video 6). Yellow cells mark future dying cells within clones (RFP, magenta). Cell membranes are labelled with *ubi-Ecad::GFP* (green). Midline regions are outlined by white dashed lines at 18h APF. Peripheral regions analysed are delimited by the aDC and pDC bristles (white peripheral lines). All cells analysed are circled at the onset of the movie. Scale bars: 10 µm.

## References

1. Aegerter-Wilmsen, T. et al. (2012) ‘Integrating force-sensing and signaling pathways in a model for the regulation of wing imaginal disc size’, Development, 139(17), pp. 3221–3231. doi: 10.1242/dev.082800.

2. Bao, S. (2014) ‘Cell adhesion in the assembly of the drosophila eye’, Journal of Neurogenetics, 28(3–4), pp. 282–290. doi: 10.3109/01677063.2014.907799.

3. Bao, S. and Cagan, R. (2005) ‘Preferential adhesion mediated by hibris and roughest regulates morphogenesis and patterning in the drosophila eye’, Developmental Cell, 8(6), pp. 925–935. doi: 10.1016/j.devcel.2005.03.011.

4. Ben-Zvi, D. S. and Volk, T. (2019) ‘Escort cell encapsulation of Drosophila germline cells is maintained by irre cell recognition module proteins’, Biology Open, 8(3), pp. 1–9. doi: 10.1242/bio.039842.

5. Bove, A. et al. (2017) ‘Local cellular neighborhood controls proliferation in cell competition’, Molecular Biology of the Cell, 28(23), pp. 3215–3228. doi: 10.1091/mbc.E17-06-0368.

6. Brás-Pereira, C. and Moreno, E. (2018) ‘Mechanical cell competition’, Current Opinion in Cell Biology, 51, pp. 15–21. doi: 10.1016/j.ceb.2017.10.003.

7. Cachoux, V. M. L. et al. (2023) ‘Epithelial apoptotic pattern emerges from global and local regulation by cell apical area’, Current Biology, 33(22), pp. 4807–4826. doi: 10.1016/j.cub.2023.09.049.

8. Cagan, R. L. and Ready, D. F. (1989) ‘The emergence of order in the Drosophila pupal retina’, Developmental Biology, 136(2), pp. 346–362. doi: 10.1016/0012-1606(89)90261-3.

9. Crozet, F. and Levayer, R. (2023) ‘Emerging roles and mechanisms of ERK pathway mechanosensing’, Cellular and Molecular Life Sciences, 80(12). doi: 10.1007/s00018-023-05007-z.

10. Donoviel, D. B. et al. (2001) ‘Proteinuria and Perinatal Lethality in Mice Lacking NEPH1, a Novel Protein with Homology to NEPHRIN’, Molecular and Cellular Biology, 21(14), pp. 4829–4836. doi: 10.1128/mcb.21.14.4829-4836.2001.

11. Dworak, H. A. et al. (2001) ‘Characterization of Drosophila hibris, a gene related to human nephrin’, Development, 128(21), pp. 4265–4276. doi: 10.1242/dev.128.21.4265.

12. Eder, D., Aegerter, C. and Basler, K. (2017) ‘Forces controlling organ growth and size’, Mechanisms of Development, 144, pp. 53–61. doi: 10.1016/j.mod.2016.11.005.

13. Eisenhoffer, G. T. et al. (2012) ‘Crowding induces live cell extrusion to maintain homeostatic cell numbers in epithelia’, Nature, 484(7395), pp. 546–549. doi: 10.1038/nature10999.

14. Grzeschik, N. A. and Knust, E. (2005) ‘IrreC/rst-mediated cell sorting during Drosophila pupal eye development depends on proper localisation of DE-cadherin’, Development, 132(9), pp. 2035–2045. doi: 10.1242/dev.01800.

15. Khoshnoodi, J. et al. (2003) ‘Nephrin Promotes Cell-Cell Adhesion through Homophilic Interactions’, American Journal of Pathology, 163(6), pp. 2337–2346. doi: 10.1016/S0002-9440(10)63590-0.

16. Lecuit, T. and Lenne, P. F. (2007) ‘Cell surface mechanics and the control of cell shape, tissue patterns and morphogenesis’, Nature Reviews Molecular Cell Biology, 8(8), pp. 633–644. doi: 10.1038/nrm2222.

17. Levayer, R., Dupont, C. and Moreno, E. (2016) ‘Tissue Crowding Induces Caspase-Dependent Competition for Space’, Current Biology, 26(5), pp. 670– 677. doi: 10.1016/j.cub.2015.12.072.

18. Linneweber, G. A., Winking, M. and Fischbach, K. F. (2015) ‘The cell adhesion molecules Roughest, Hibris, Kin of irre and Sticks and Stones are required for long range spacing of the Drosophila wing disc sensory sensilla’, PLoS ONE, 10(6), pp. 1–21. doi: 10.1371/journal.pone.0128490.

19. Marinari, E. et al. (2012) ‘Live-cell delamination counterbalances epithelial growth to limit tissue overcrowding’, Nature, 484(7395), pp. 542–545. doi: 10.1038/nature10984.

20. Matamoro-Vidal, A. and Levayer, R. (2019) ‘Multiple Influences of Mechanical Forces on Cell Competition’, Current Biology, 29(15), pp. R762–R774. doi: 10.1016/j.cub.2019.06.030.

21. Merino, M. M. et al. (2015) ‘Elimination of unfit cells maintains tissue health and prolongs lifespan’, Cell, 160(3), pp. 461–476. doi: 10.1016/j.cell.2014.12.017.

22. Moreno, E. et al. (2019) ‘Competition for Space Induces Cell Elimination through Compaction-Driven ERK Downregulation’, Current Biology, 29(1), pp. 23–34.e8. doi: 10.1016/j.cub.2018.11.007.

23. Mummery-Widmer, J. L. et al. (2009) ‘Genome-wide analysis of Notch signalling in Drosophila by transgenic RNAi’, Nature, 458(7241), pp. 987–992. doi: 10.1038/nature07936.

24. Nagarkar-Jaiswal, S. et al. (2015) ‘A library of MiMICs allows tagging of genes and reversible, spatial and temporal knockdown of proteins in Drosophila’, eLife, 4(3), pp. 1–28. doi: 10.7554/eLife.05338.

25. Paci, G. and Mao, Y. (2021) ‘Forced into shape: Mechanical forces in Drosophila development and homeostasis’, Seminars in Cell and Developmental Biology, 120(March), pp. 160–170. doi: 10.1016/j.semcdb.2021.05.026.

26. Paul, A. et al. (2022) ‘Cell adhesion molecule KIRREL1 is a feedback regulator of Hippo signaling recruiting SAV1 to cell-cell contact sites’, Nature Communications, 13(1), pp. 1–14. doi: 10.1038/s41467-022-28567-3.

27. Pereira, A. M. et al. (2011) ‘Integrin-dependent activation of the jnk signaling pathway by mechanical stress’, PLoS ONE, 6(12). doi: 10.1371/journal.pone.0026182.

28. Rhiner, C. et al. (2010) ‘Flower forms an extracellular code that reveals the fitness of a cell to its neighbors in Drosophila’, Developmental Cell, 18(6), pp. 985–998. doi: 10.1016/j.devcel.2010.05.010.

29. Shraiman, B. I. (2005) ‘Mechanical feedback as a possible regulator of tissue growth’, PNAS, 102(9), pp. 3318–3323. doi: 10.1073/pnas.0404782102.

30. Simons, M., Hartleben, B. and Huber, T. B. (2009) ‘Podocyte polarity signalling’, Current Opinion in Nephrology and Hypertension, 18(4), pp. 324–330. doi: 10.1097/MNH.0b013e32832e316d.

31. Takemura, M. and Adachi-Yamada, T. (2011) ‘Cell death and selective adhesion reorganize the dorsoventral boundary for zigzag patterning of Drosophila wing margin hairs’, Developmental Biology, 357(2), pp. 336–346. doi: 10.1016/j.ydbio.2011.07.007.

32. Tryggvason, K., Patrakka, J. and Wartiovaara, J. (2006) ‘Hereditary Proteinuria Syndromes and Mechanisms of Proteinuria’, New England Journal of Medicine, 354(13), pp. 1387–1401. doi: 10.1056/nejmra052131.

33. Tsuboi, A. et al. (2017) ‘Inference of cell mechanics in heterogeneous epithelial tissue based on multivariate clone shape quantification’, Frontiers in Cell and Developmental Biology, 5(AUG). doi: 10.3389/fcell.2017.00068.

34. Valon, L. et al. (2021) ‘Robustness of epithelial sealing is an emerging property of local ERK feedback driven by cell elimination’, Developmental Cell, 56(12), pp. 1700–1711.e8. doi: 10.1016/j.devcel.2021.05.006.

35. Villars, A. et al. (2022) ‘Microtubule disassembly by caspases is an important rate-limiting step of cell extrusion’, Nature Communications, 13(1). doi: 10.1038/s41467-022-31266-8.

36. Wagstaff, L. et al. (2016) ‘Mechanical cell competition kills cells via induction of lethal p53 levels’, Nature Communications, 7. doi: 10.1038/ncomms11373.

37. Yamamoto, M. et al. (2017) ‘The ligand Sas and its receptor PTP10D drive tumour-suppressive cell competition’, Nature, 542(7640), pp. 246–250. doi: 10.1038/nature21033.

